# Application of long-read sequencing for robust identification of correct alleles in genome edited animals

**DOI:** 10.1101/838193

**Authors:** Christopher V. McCabe, Gemma F. Codner, Alasdair J. Allan, Adam Caulder, Skevoulla Christou, Jorik Loeffler, Matthew Mackenzie, Elke Malzer, Joffrey Mianné, Fran J. Pike, Marie Hutchison, Michelle E. Stewart, Hilary Gates, Sara Wells, Nicholas D. Sanderson, Lydia Teboul

## Abstract

Recent developments in CRISPR/Cas9 genome editing tools have facilitated the introduction of more complex alleles, often spanning genetic intervals of several kilobases, directly into the embryo. These techniques often produce mosaic founder animals and the introduction of donor templates, via homologous directed repair, can be erroneous or incomplete. Newly generated alleles must be verified at the sequence level across the targeted locus. Screening for the presence of the desired mutant allele using traditional sequencing methods can be challenging due to the size of the desired edit(s) together with founder mosaicism. In order to help disentangle the genetic complexity of these animals, we tested the application of Oxford Nanopore long read sequencing of the targeted locus. Taking advantage of sequencing the entire length of the segment in each single read, we were able to determine whether the entire intended mutant sequence was present in both mosaic founders and their offspring.

## Background

Genome editing tools in conjunction with a single-stranded oligonucleotide (ssODN) donor are an effective method for the introduction of specific mutations in early embryos (Wang et al., 2013; Yang et al., 2013). However, this strategy often produces complex mosaic animals in the founder (G_0_) generation (Singh et al., 2015; Mianné et al., 2017). Each of the founder alleles with evidence of the desired edits must be fully sequenced in order to detect unwanted mutations *in cis* (Mianné et al., 2016; Renaud et al., 2016; Birling et al., 2017).

Previously, Sanger sequencing has been sufficient to characterise CRISPR/Cas9 mutagenised loci using ssODN donors. These templates were a maximum of 200 bases in length (Mianné et al., 2017), which can be easily covered within a Sanger sequencing read. However, long single-stranded DNA (lssDNA) donors (Quadros et al., 2017; Codner et al., 2018) or multiple ssODNs (Yang et al., 2013; Lanza et al., 2018) can be used for the generation of complex alleles directly in one-cell embryos. Targeted edits spanning several kilobases (kb) are now being produced with increasing regularity. In order to cover intervals of this size, and piece together each of the many allele variants generated in a mosaic founder, several 500-to 800 bp Sanger sequencing reads are required and subsequently combined *in silico* (Figure 1 and Supplemental Figure 1, process highlighted in orange). As such, the characterisation of mutant alleles in mosaic founders is particularly challenging. Events that are not captured by the chosen assays can be omitted (Shin et al., 2017; Kosicki et al., 2018; Owens et al., 2019) and screening can sometimes fail to distinguish rearranged from correct alleles in these complex animals (Codner et al., 2018). This suggests that the screen based on Sanger sequencing produced some false positives.

**Figure 1:**
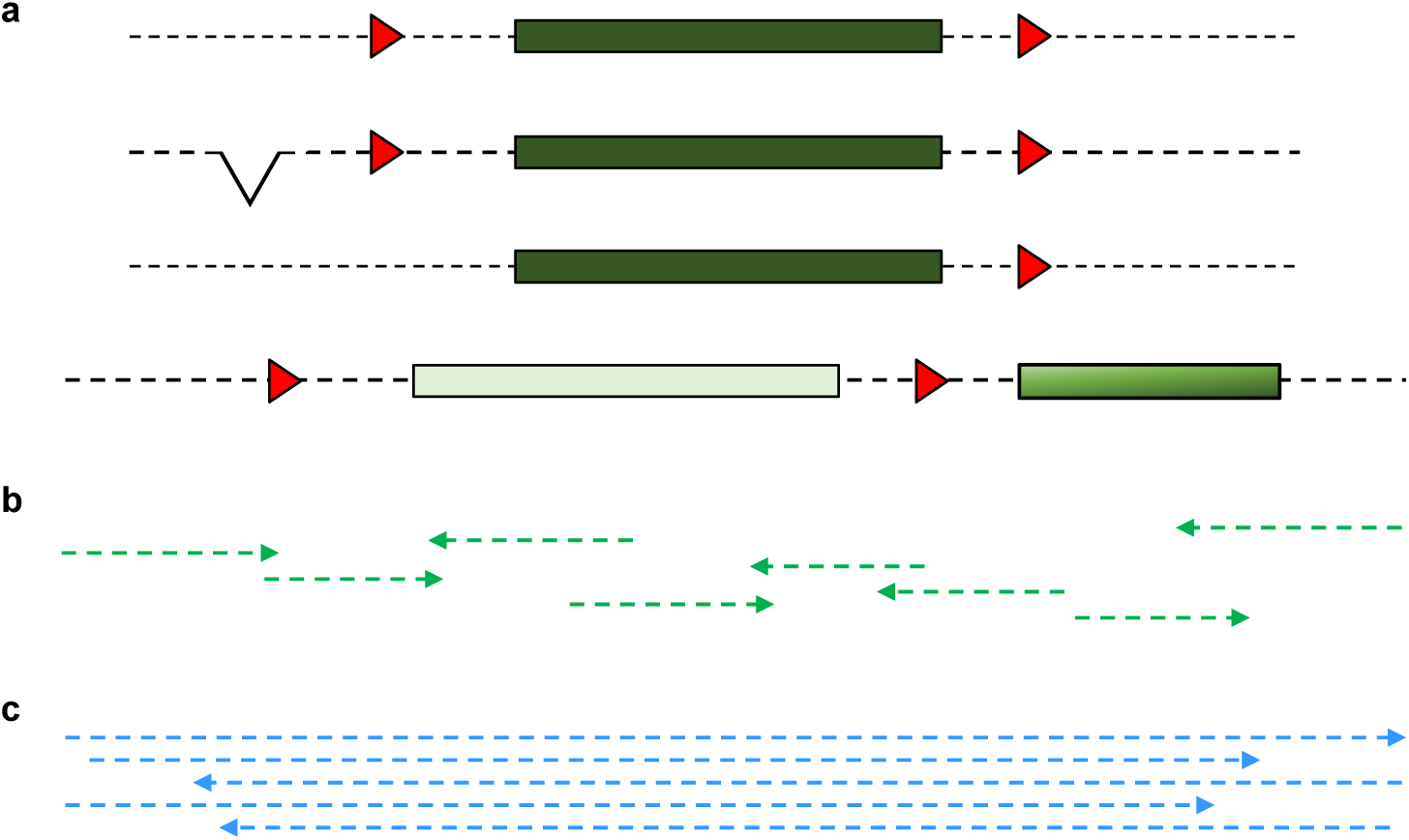
Disentangling sequences from a mosaic animal. (a) illustrates the complex genetic make up at a targeted locus in a mosaic (four alleles). Red triangles correspond to loxP sites, green boxes indicate the floxed exon. The mosaic animal contains: correct floxed allele, floxed allele with a deletion upstream, a partial integration of the donor (3’ LoxP only) and a donor integration with a duplicated segment. (b) illustrates how several Sanger sequencing reads are required to span the whole interval interest. Note that it is impossible to ascertain how reads should be assembled to span each specific allele. (c) illustrates how each long read spans the whole interval of interest sequenced within a mosaic.

Oxford Nanopore Technologies (ONT) sequencing produces far longer reads, which can easily cover the entire length of the mutagenised interval in one molecule (Jain et al., 2016). We piloted the use of ONT sequencing as an alternative method for identifying the presence of the correctly mutated allele in mosaic founders, derived from the microinjection of CRISPR/Cas9 reagents and lssDNA donors, and their progeny (G_1_; Figure 1 and process highlighted in blue in Supplemental Figure 1). We showed that the high error rate inherent to ONT sequencing can be offset by very deep sequencing coverage. We assembled a workflow for the analysis of sequencing data to identify the genomic DNA samples that include the correct allele. We found that ONT sequencing provides an accurate screen of these animals.

Importantly, long reads allow for the earlier exclusion of founder animals that were identified as positive for the presence of a correct integration by Sanger sequencing based screen but only transmitted incorrectly mutated alleles. The application of ONT sequencing for the screening of founders obtained with lssDNA donors represents an advance for ethical animal use, as it prevents breeding of some false-positive founders.

## Results

### Establishing an accurate ONT-based targeted sequencing screening process

As ONT has a higher error rate than other next generation sequencing technologies (Jain et al., 2016), we first assessed the feasibility of unequivocally recognising known sequences and defined which quality thresholds can be used for such analysis. We analysed sequencing data from six PCR amplicons amplified from WT animal biopsies with tailed primers flanking genomic intervals ranging from 0.9 to 2 kb in size (Experiment A, Supplemental Table ST1). PCR amplicons were barcoded, assembled in sequencing libraries and sequenced with a MinION.

Using the Porechop tool, reads that showed both barcoded ends were selected and demultiplexed for each barcode. We then filtered reads according to a range of read quality filters (q84 to q96) with Filtlong. Each group of reads of a given quality threshold was aligned against the genome reference sequence using Minimap2 (Li, 2018). We then evaluated targeted sequencing performance by its ability to make a call for each base in the interval (coverage breadth) and the sequencing accuracy of these calls at the base level.

As expected, higher quality filters retained fewer reads (Supplemental Figure 2a), with the number of reads dropping sharply from q95. At the highest quality thresholds, although the reads were of superior quality, there were not enough of them to achieve complete coverage over the target interval (Supplemental Figure 2b). This is in contrast to the larger numbers of reads retained at less stringent quality filters, that did achieve complete coverage over the target interval. Intermediate read depths (100X) identify very high proportions of the genome reference sequences. With a quality filter set between q86 and q94, when very high depth of reads (1,000X-10,000X) remained after filtering, the process was able to cover all bases within the sequenced segments.

**Figure 2:**
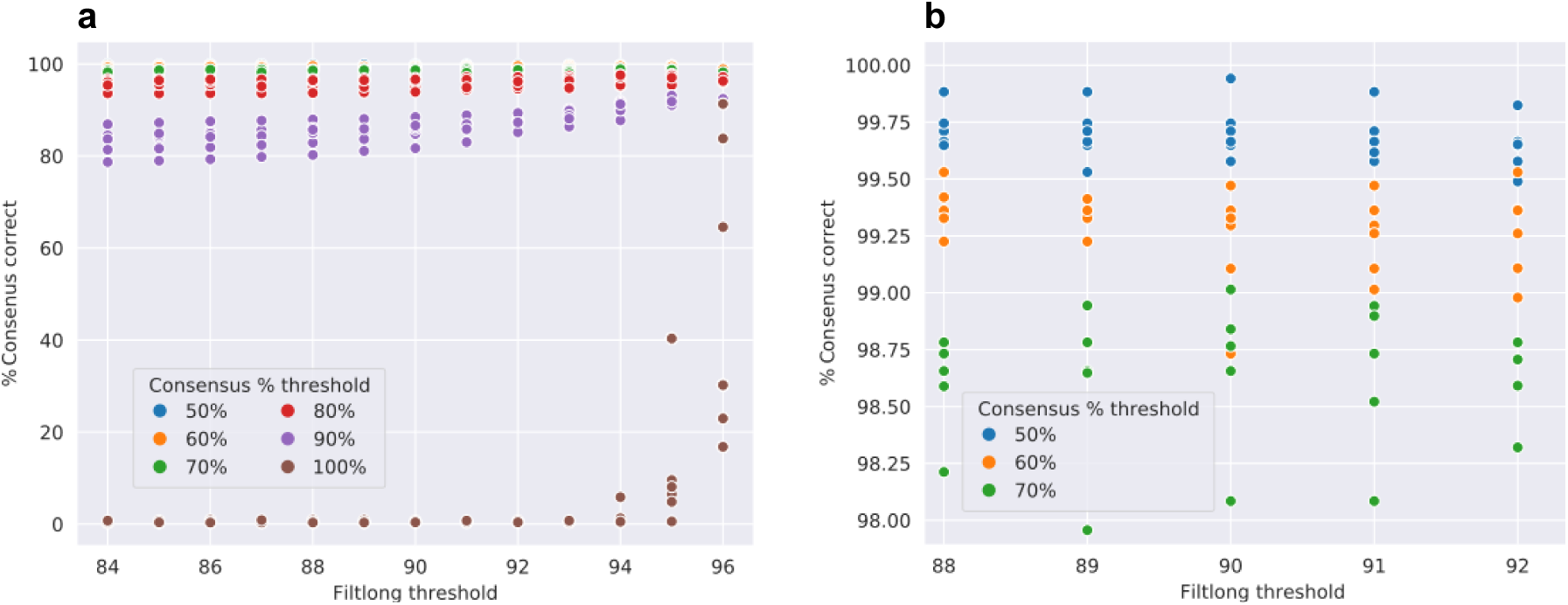
The graphs show the percentage of bases accurately sequenced by ONT sequencing for each WT segment analysed, for Filtlong read quality filters from q90 to q96 and with base calling threshold between 50% to 100% with increments of 10%. (a) Large numbers of calls achieved full coverage and highest accuracy for each base. (b) shows a zoomed in portion of (a). Very high percentages of sequence identification (>99.5%) are achieved with intermediate Filtlong quality threshold (q90) and lower threshold for consensus base calling (50%).

We then analysed sequencing accuracy with thresholds for base-calling (proportion of calls at each base position that has to be reached to declare a consensus call for the base) ranging from 50% to 100%. Figure 2 shows the percentage of reference sequence accurately identified with consensus thresholds ranging from 50% to 100%, for each of the six loci. Figure 2a shows that selecting the highest quality read did not achieve the highest sequencing accuracy. Figure 2b focuses on the parameter ranges that ensure a very high proportion of accurate sequence identification. The graph shows that in our experimental conditions q90 Filtlong read quality threshold and a 50% consensus base calling threshold supports >99.5% sequence accuracy. We therefore selected these parameters for the remainder of the study.

### ONT-based sequencing analysis of mutants generated with CRISPR/Cas9 and lssDNA donors

In the next run we analysed founder animals and their offspring (Experiment B, Supplemental Table ST1) for two projects: a Cre KI into the *Mpeg1* gene, (Figure 3a) and a floxed *Cx3cl1* allele (Figure 3b). The sequence of donor lssDNAs and primers used in this article is shown in Supplemental Table ST2 and the animals produced in this study are summarised in Supplemental Table ST3. All of these animals had been previously identified as potentially bearing the desired allele change by Sanger sequencing (Supplemental Figures S3 and S4). PCR products amplified from both G_0_ and G_1_ animals with external primers were sequenced with ONT sequencing. Q90 reads were aligned to the intended mutant sequence. Alignments were visualised with IGV (Thorvaldsdóttir et al., 2013; Figure 4). The sequence of the Mpeg1-cre allele was fully confirmed in both the positive founder Mpeg1-cre-80 and their offspring (Figure 4a).

**Figure 3:**
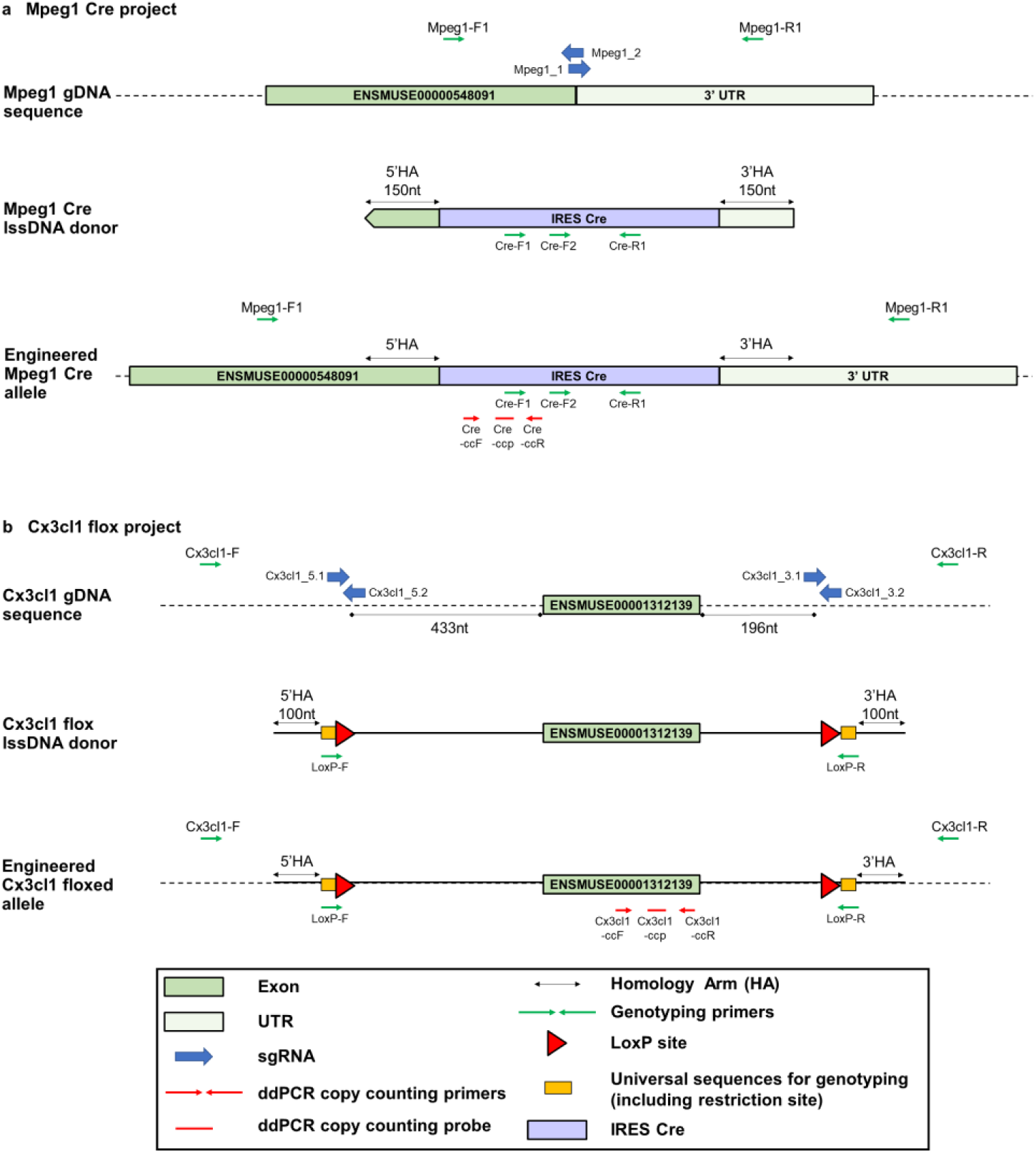
The figure details the design of the (a) Mpeg1-cre and the (b) Cx3cl1-flox (Codner et al., 2018) allele, respectively. The position of the primers used for analysis (sequences detailed in Supplemental Table 2) are shown, together with that of the ddPCR copy counting assays.

**Figure 4:**
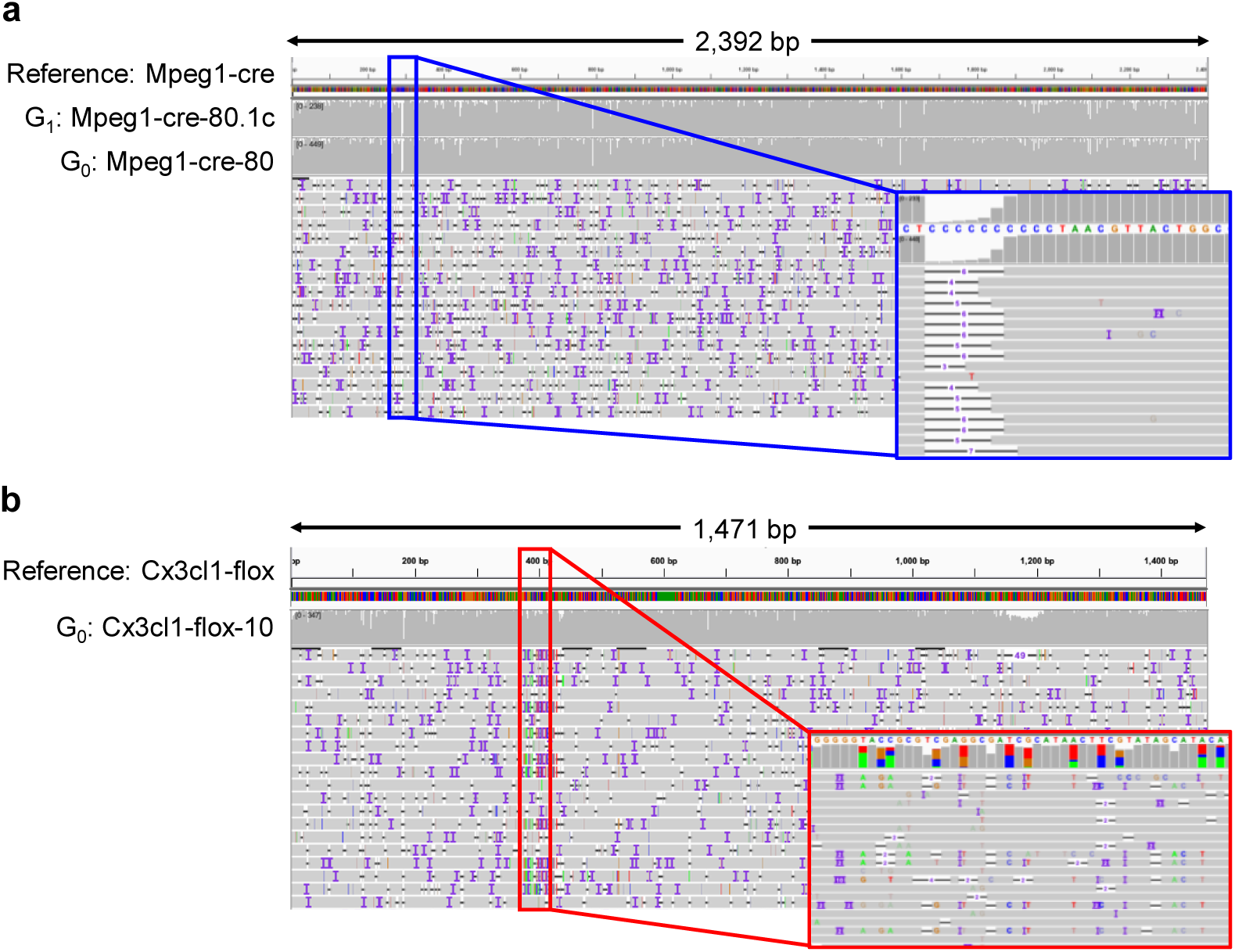
The figure summarises the outcome of ONT sequencing of animals for (a) the Mpeg1-cre and (b) Cx3cl1-flox project vizualised with IGV. The alignment reflects the noisy nature of the method with errors distributed across the length of the sequenced segment. Note the complete alignment of reads to designed mutant sequences for both mosaic founders (G_0_s) and G_1_ (grey histograms). Note that the different alleles in mosaic founder. Some reads align to the reference across the whole interval (red frame). The dip in sequence coverage coincides with at homopolymer repeat (Mpeg1-cre-80 and Mpeg1-cre-80.1c, blue frames).

### Aiding the identification of mutant alleles

For floxed alleles, the WT and mutant sequences only differ by a small proportion of their overall length (typically 120 bases out of 1.5 kb which is less than ONT sequencing raw read error frequency). Consequently, a high stringency parameter for alignment is not sufficient to prevent WT reads from aligning against the mutant reference along with mutant reads, potentially producing an ambiguous alignment file. To refine the analysis, we filtered reads for the presence of segments exclusive to the mutant sequence (determinant) prior to generating alignments. Typically two determinants for floxed alleles, or one determinant for cassette KIs can be used. This yielded unequivocal sequence alignments and the correct mutated sequence was detected at the founder stage (Figure 4b).

Founder animals from four more floxed allele projects were tested in this run using PCR amplicons generated using tailed-primers external to the donor templates (generic design shown in Supplemental Figure S5, Project Prdm8-flox, Pam-flox, Hnf1a-flox and Inpp5k-flox; summary of samples in Experiment B, Supplementary Table ST1 and Sanger sequencing-based characterisation of mice in Supplemental Figures S6, S7, S8 and S9 respectively). ONT sequencing showed that the PCR amplicons amplified from founders Prdm8-flox-31, Pam-flox-3 and Hnf1a-flox-66 bear the correct sequences (Figure 5a and Supplemental Figure S10, respectively). Prdm8-flox-31 and Hnf1a-flox-66 matings had yet to produce G_1_ animals but PCR amplicons amplified from Pam-flox-3 offspring were sequenced and confirmed as correct (Supplemental Figure S7).

**Figure 5:**
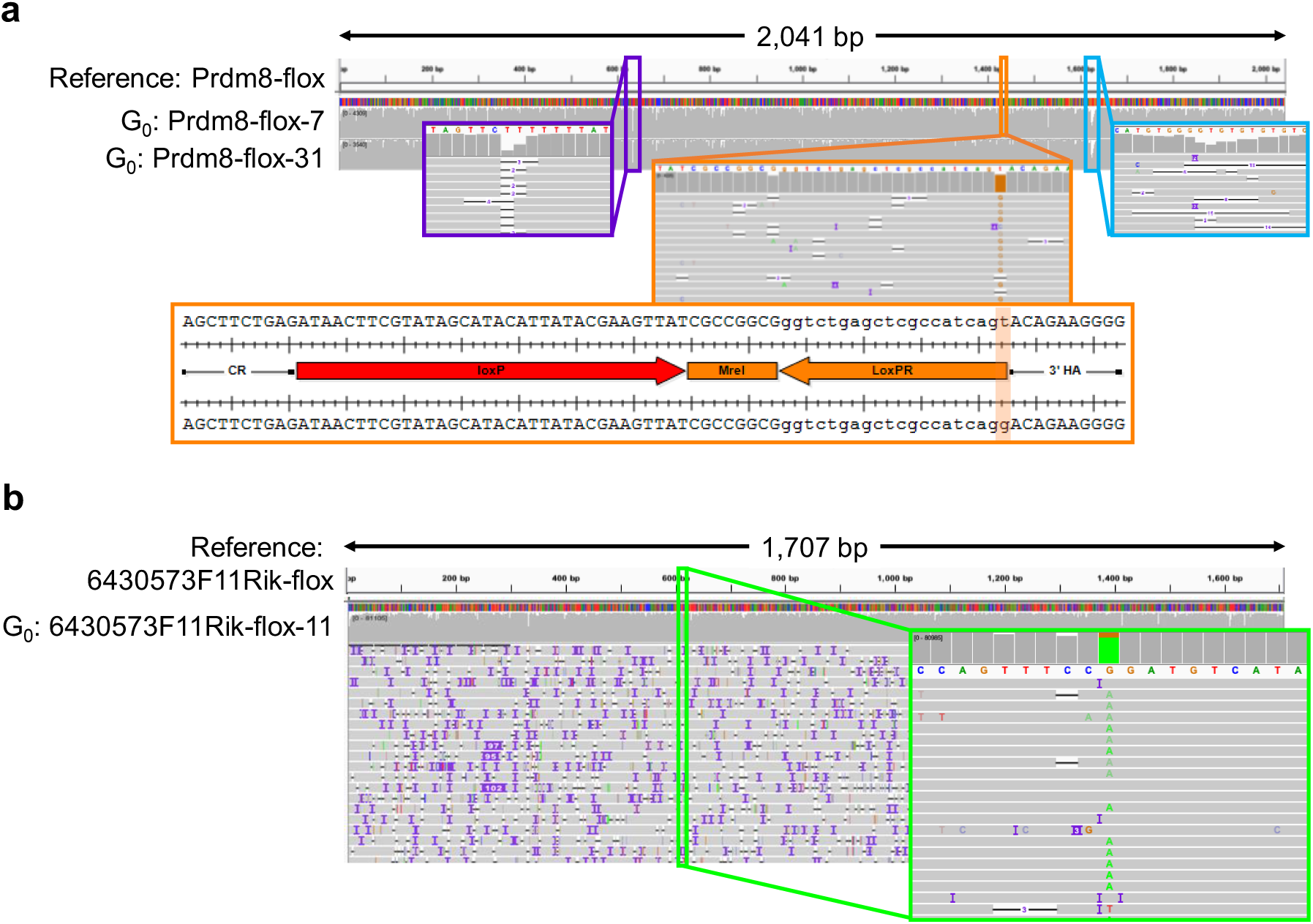
The figure shows alignments of sequencing reads obtained from founders Prdm8-flox-7 and Prdm8-flox-31 (a) and 6430573F11Rik-flox-11 (b) against the designed respective floxed sequence. ONT sequencing reveals an *in cis* point mutation, not present in the donor, associated with alleles that contained loxP sites in both Prdm8-flox-7 (a, orange frames) and 6430573F11Rik-flox-11 (b, green frames). The same sequences were confirmed with Sanger reads in these animals (Codner et al., 2018 and this article, Supplemental Figures S7 and S10). Note that dips in sequence coverage coincide with repeated sequences (Prdm8-flox-7 and Prdm8-flox-31, purple and blue frames, Figure 5a).

Sanger sequencing of Inpp5k-flox allele founders showed the presence of loxP sites but did not lead to a conclusive outcome due to the mosaic nature of the template (Supplemental Figure S9). ONT sequencing of a PCR amplicon obtained from founder Inpp5k-flox-7 without preliminary filtering of reads for the presence of both loxPs revealed the genetic complexity of the animals (Supplemental Figure S11). Importantly, filtering for the presence of both loxP determinants resulted in no reads aligning to the designed mutant sequence. This absence of the fully conforming allele is in keeping with the result of Sanger sequencing the offspring of founder Inpp5k-flox-7 (Supplemental Figure S12).

### Unwanted Single Nucleotide Polymorphisms (SNPs) can be visualised in ONT alignments

Having passed filtering with determinants, PCR amplicons amplified from Prdm8-flox-7 showed the mutant allele contained an unintended point mutation associated with the flox sequences (Figure 5a). The point mutation is in the synthetic interval flanking the 3’ loxP and will not affect future use of this new allele. The SNP was also identified in Sanger sequencing of this individual (Supplemental Figure S6), and in the subsequent generation (data not shown).

In a third ONT sequencing run (Experiment C, Supplemental Table ST1), visualisation of the alignment file highlighted that mosaic founder 6430573F11Rik-flox-11 showed a G to A change at position 616 present at high representation (Figure 5b). This unintended point mutation was systematically associated *in cis* with alleles that contained loxP sites. These mutations are also seen in Sanger read data from this individual (Supplemental Figure S13).

## Discussion

### Unequivocal identification of positive founders

Here we demonstrate that ONT, in spite of a higher per-base error rate, can be employed to efficiently identify correctly targeted alleles when screening mosaic G_0_s (Figures 4 and 5). This can be extended to G_1_ to validate the transmitted allele and to confirm segments difficult to sequence by standard sequencing, such as regions downstream of homopolymer repeats (Figure 4).

### Depth of sequencing offsets sequencing error rate

We have optimised our strategy for sequencing and data analysis workflow by sequencing targeted regions in WT animals. Importantly, confidence in sequencing data was achieved from the extensive depth of coverage, rather than through setting the most stringent quality filters for sequencing data (Figure 2). With sufficient sequencing read depth, reads were mapped to WT reference sequences concordantly across the entire genomic interval (Experiment A). The depth of coverage required depended on the complexity of the sequence with repeated sequences requiring deeper sequencing (Figures 4a and 5a). We then interrogated the sequencing data obtained from mutant animals for multiple projects for previously verified mutant alleles and found they could be unequivocally detected (Experiment B; Projects Mpeg1-cre, Cx3cl1-flox, Prdm8-flox and Pam-flox). In all cases where WT and mutant sequences differ by a proportion smaller than or close to the sequencing error rate, it was preferable to filter reads for the presence of determinants specific to mutant reference sequences prior to alignment. This prevents reads that correspond to WT alleles or partial integrations of mutant donors from being included in the alignments. This produced an unambiguous readout for the presence of correct mutant alleles. Homopolymer repeats were the exception to this conclusion as such segments remain a challenge for any sequencing techniques.

### Exclusion of Sanger sequencing-based false positive animals

We applied ONT sequencing to projects where the presence of desired sequences had been shown in some founder G_0_ animals employing Sanger sequencing but only imperfectly mutated alleles were found transmitted to the subsequent generation (Projects 6430573F11Rik-flox, Figure 5b and Inpp5k-flox, Supplemental Figure S13). Crucially, application of ONT sequencing to these same founders showed that the desired mutation was systematically associated with additional base-pair changes or deletion *in cis*. Therefore, a long-read sequencing-based screening strategy can support the identification of undesired mutant alleles at an earlier stage of the mutagenesis process (G_0_ screening). By contrast, with a screening strategy based on Sanger sequencing, the definitive sequence of each allele can only been ascertained in the simpler, non-mosaic, G_1_s. Importantly, the ONT approach is also applicable when a pair of ssODN donors is employed instead of a lssDNA (Yang et al., 2013), allowing the identification of animals where the two sequences have been correctly integrated in *cis*.

### Run capacity

We have used targeted sequencing of relatively short PCR amplicons (up to approximately 3 kb) permiting the generation of ultra-deep (1,000X to 10,000X) coverage datasets. This uses a fraction of the sequencing data production capacity of a MinION run. We used a kit that supports twelve barcodes so that twelve animals can be sequenced in parallel analysing the same genomic segment. It would be possible to further multiplex samples by designing alternative primer pairs to amplify the same core region of interest but which include different flanking region lengths. Differentiation between individual animals for the same locus/project within a run can then be achieved using the genomic context flanking the region as an internal barcode. With the alternative format of 96 barcodes, a conservative set-up of 96 individuals (each under one barcode), if they were all mosaic, would require in the order of twenty gigabases sequenced to achieve 10,000X coverage of a 3 kb segment (96 animals for a given project containing up to eight genetic identities), which can be produced in a single MinION run.

We note that close to full identification of the target region was obtained with much lower coverage, offering the possibility of screening many more samples within a run with very high reliability. Finally, we have used an additional dimension for multiplexing, as animals corresponding to two different projects can be analysed in parallel under the same barcode. More samples could have been multiplexed using this strategy.

Large numbers of reads, rather than read quality, underpin the accurate recognition of the desired mutant sequences. However, it is noteworthy that all sequencing runs we employed for the study were interrupted within twenty four hours, well before the standard forty eight hours recommended by the manufacturer, to reduce the amount of sequencing data excessive to our purpose being generated.

Currently, the main limiting factors of this process are the reliance on PCR, which may introduce sequence errors and the length of PCR product that can be amplified from genomic DNA extracted from a tissue biopsy. The recently proposed nanopore Cas9 Targeted-sequencing (nCATS) facilitates targeted sequencing by ONT without PCR and may increase the size of the genomic segments that can be surveyed for more extensive validation of animals (Gilpatrick et al., 2019).

### A simple process

Although the approach appears a step change at first glance, the use of long-read sequencing turns out to involve a fairly simple and accessible process. It requires only minimal investment in sequencing equipment. The generation of sufficient sequencing data utilises a small fraction of the system capacity and the datasets that are produced do not represent a challenge for existing analysis tools. We further facilitate access to the process by packaging the analysis in a simple workflow (see Methods). The timeline from genomic DNA extraction from a potential founder to a fully analysed dataset informing on the presence of the desired mutant allele fits within one week. ONT sequencing is a simple and efficient tool for screening genome-edited founders obtained with lssDNA donors. This is in contrast to traditional Sanger sequencing methods which rely on the amplification of multiple PCR products, which must then be individually sequenced and assembled into contigs. Assembly of these contigs is liable to mis-associate *trans* reads in mosaics animals, resulting in false positive calls.

However, the extensive characterisation of the target loci that ONT sequencing supports, in both founder and subsequent generations, does not suffice to validate newly mutated lines. Indeed, checks of off-target events (Anderson et al., 2018; Iyer et al., 2018), in particular, those physically linked to the locus of interest) and copy counting of the donor sequence by ddPCR to eliminate additional integrations remain essential and complementary steps to fully validate G_1_ animals (Supplemental Figures S3 and S4 and Codner et al., 2018).

Finally, this simple sequencing process is also applicable to any other circumstance where a sequence of a specific locus must be validated, for example in cultured cells following gene targeting by homologous recombination.

### A more accurate screening tool for more ethical animal management

Here we have illustrated how long-read sequencing can be employed to exclude founders previously misidentified as positive using Sanger sequencing-based methods, because they only contained targeted mutations that were associated with unwanted base-pair changes or sequence rearrangements. The method also allows for the analysis of mosaic animals where the genetic make-up is too complex to be easily disentangled by standard Sanger sequencing. This constitutes a refinement in terms of the use of animals for the generation of targeted mutations, as it reduces the number of false positive founders carried forward for breeding. It also serves to shorten the timeline of mutagenesis projects, as founder characterisation is greatly facilitated and no animals nor time are unnecessarily used testing misidentified positive founders for germline transmission of the desired mutant allele. This is particularly useful as founders can present a broad range of welfare issues, as part or all of their body may contain mutations that may affect both alleles of modified loci.

## Conclusion

CRISPR/Cas9 with lssDNA donors delivered into one-cell embryos generates complex mosaic founders that are challenging to analyse by classical Sanger sequencing. We showed that targeted sequencing with ONT technology is a simple and powerful method to faithfully identify the animals that bear a correct integration on target. This represents progress in ethical animal use, as it prevents breeding of false-positive founders.

## Materials and methods

### Sequences of reagents

The sequences of the sgRNAs, templates for lssDNA generation, primers and probes are shown in Supplemental Table ST2.

### sgRNAs

Guide sequence selection was carried out using the following online tools: CRISPOR (Haeussler et al., 2016) and WTSI Genome Editing (WGE) (Hodgkins et al., 2015). sgRNA sequences were selected with as few predicted off-target events as possible, particularly on the same chromosome as the intended modification. sgRNAs used in this study are shown in Table ST2. sgRNAs were synthesised directly from gBlock^®^ (IDT) templates containing the T7 promoter using the HiScribe™ T7 high yield RNA synthesis kit (New England BioLabs^®^) following manufacturer’s instructions. RNAs were purified using the MEGAclear kit (Ambion). RNA quality was assessed using a NanoDrop (ThermoScientific) and by electrophoresis on 2% agarose gel containing ethidium bromide (Fisher Scientific).

### Templates for lssDNA synthesis

Templates for lssDNA synthesis were either assembled by cloning in a plasmid or, when possible, were obtained from IDT as a single gBlock^®^.

### Donor templates

Donor lssDNAs were generated following a method adapted from Miura et al., 2015. Briefly, templates for *in vitro* transcription (donor sequence flanked by the T7 promoter) were obtained as a gBlock^®^ (IDT) or cloned in a plasmid that was subsequently linearised. Typically, 150 ng of double stranded gBlock^®^ template or 2 µg of plasmid template was transcribed using the HiScribe T7 High Yield RNA Synthesis Kit (New England BioLabs^®^). At the end of the reaction, DNase I was added to remove the DNA template. RNA was purified employing the MEGAclear Transcription Clean-Up kit (Ambion). Single-stranded DNA was synthesised by reverse transcription from 20 µg of RNA template employing SuperScript III Reverse Transcriptase (Invitrogen), treated with RNAse H (Ambion) and purified employing the QIAquick Gel Extraction Kit (Qiagen) or, for higher yields, employing the RNA Clean & Concentrator™ kit (Zymogen). Alternatively, lssDNAs were synthetised with the Guide-it™ Long ssDNA Strandase Kit according to the manufacturer instruction. Donor concentration was quantified using a NanoDrop (Thermo Scientific) and integrity was checked on 1.5% agarose gel containing ethidium bromide (Fisher Scientific).

### Mixes for microinjection

Microinjection buffer (10 mM Tris-HCl, 0.1 mM EDTA, 100 mM NaCl, pH7.5) was prepared and filtered through a 2 nm filter and autoclaved. Mixes containing 100 ng/µl Cas9 mRNA (5meC,Ψ) (TriLink BioTechnologies), 50 ng/µl sgRNAs and 50 ng/µl ssODN or 50 ng/µl lssDNA were prepared in microinjection buffer, filtered through Costar^®^ SpinX^®^ Centrifuge Tube Filters (Corning) and stored at -80°C until microinjection.

### Mice

All animals were housed and maintained in the Mary Lyon Centre, MRC Harwell Institute under specific pathogen-free (SPF) conditions, in individually ventilated cages adhering to environmental conditions as outlined in the Home Office Code of Practice. Mice were euthanised by Home Office Schedule 1 methods. Animals used for transgenesis projects are detailed in Supplemental Table ST3. Colonies established during the course of this study are available for distribution and are detailed in Supplemental Table ST4.

### Pronuclear microinjection of zygotes

All embryos were obtained by superovulation. Pronuclear microinjection was performed as per Gardiner and Teboul, 2009, employing a FemtoJet (Eppendorf) and C57BL/6NTac embryos. Specifically, injection pressure (Pi) was set between 100 and 700 hPa, depending on needle opening; injection time (Ti) was set at 0.5 seconds and the compensation pressure (PC) was set at 10 hPa. Mixes were centrifuged at high speed for one minute prior to microinjection. Injected embryos were re-implanted in CD-1 pseudo-pregnant females. Host females were allowed to litter and rear G_0_s.

### Breeding for germline transmission

G_0_ animals where the presence of a desired allele was detected were mated to WT isogenic animals to obtain G_1_ animals to assess the germline transmission of the allele of interest and permit the definitive validation of its integrity.

### Genomic DNA extraction from ear biopsies

Genomic DNA from G_0_ and G_1_ animals was extracted from ear clip biopsies using the DNA Extract All Reagents Kit (Applied Biosystems) according to manufacturer’s instructions. The crude lysate was stored at -20°C.

### PCR amplification and Sanger sequencing

New primer pairs were set up in a PCR reaction containing 500 ng genomic DNA extracted from a wild type mouse, 1 x Expand Long Range Buffer with 12.5 mM MgCl_2_ (Roche), 500 µM PCR Nucleotide Mix (dATP, dCTP, dGTP, dTTP at 10 mM, Roche), 0.3 µM of each primer, 3% DMSO, and 1.8 U Expand Long Range Enzyme mix (Roche) in a total volume of 25 µl. Using a T100 thermocycler (Bio-Rad), PCRs were subject to the following thermal conditions; 92°C for 2 minutes followed by 40 cycles of 92°C for 10 seconds, a gradient of annealing temperatures between 55-65°C for 15 seconds and 68°C for 1 minute/kb and a final elongation step for 10 minutes at 68°C. PCR outcome was analysed on a 1.5 to 2% agarose gel, depending on the amplicon size and the highest efficient annealing temperature was identified for the primer pair. If no temperature allowed for an efficient and/or specific PCR amplification the assay was repeated with an increased DMSO concentration (up to 12%). Using optimised conditions, as defined above, PCRs for each project were run and an aliquot analysed on agarose gel. PCR products were purified employing QIAquick Gel Extraction Kit (Qiagen) and sent for Sanger sequencing (Source Bioscience, Oxford). Genotyping primers were chosen at least at 200 bp away from the extremity of donors, depending on available sequences for design.

### Analysis of Sanger sequencing data

Sequencing data were analysed differently depending on whether they were obtained from G_0_s or G_1_s (as per Gardiner and Teboul, 2009). At the G_0_ stage, animals were screened for evidence of the expected change i.e. the presence of loxP sites for conditional allele projects or presence of the cre knock-in sequence for Mpeg-cre allele. G_0_ animals should be considered mosaic animals. All G_1_ animals are heterozygous containing one WT allele and one allele to be determined as they are obtained from mating G_0_ animals with desired gene edits to WT animals. The G_1_ stage enables definitive characterisation of the new mutant.

### Preparation of libraries for ONT sequencing

DNA LoBind tubes (Eppendorf) were used. PCR was performed with tailed-end primers using the same conditions as for amplicons produced for Sanger sequencing, to generate amplicons for ONT sequencing. PCR amplicons were barcoded employing LongAmp Taq (New England BioLabs^®^). The ends of pooled DNA fragments were repaired employing the NEBNext End repair / dA-tailing Module (New England BioLabs^®^). Sequencing adaptors were added using the 1D-Ligation Sequencing Kit (Oxford ONT technology). All reactions were performed according to the manufacturer’s instruction. DNA was purified at all steps using AMPure XP beads (Agencourt) employing a 0.8X to 1X beads to sample ratio. DNA was quantified with a Qubit fluorometer at all steps. Sequencing libraries were loaded on primed SpotON Flow Cell (R9.4) (ONT). Runs were performed employing the MinKNOW GUI at defaults settings for up to 24 hours (ONT).

### Analysis of ONT sequencing data

A nextflow (Di Tommaso et al., 2017) workflow for the bioinformatics processes was assembled and is available on gitlab (https://gitlab.com/nick297/cas9point4). In brief, reads were demultiplexed with Porechop (Wick et al., 2017), requiring the recognition of two barcodes (both extremities of the PCR amplicon sequenced). Reads were then filtered for a minimum quality (q score) using Filtlong (https://github.com/rrwick/Filtlong). Filtered reads were aligned against the relevant reference sequence using minimap2 (Li, 2018) and filtered using samtools for a mapping quality score of q90 (Li et al., 2009). Alignments were then visualised using IGV (http://software.broadinstitute.org/software/igv/; Thorvaldsdóttir et al., 2013). After alignment, corresponding reads were filtered for the presence of mutant specific sequence determinants using BLASTn (Altschul et al., 1990) to simplify the readout of the experiment.

### Copy counting of the donor by ddPCR

Copy Number Variation experiments were performed as duplex reactions. A FAM-labelled assay was used to amplify a region contained within the ssDNA donor (sourced from Biosearch Technologies), in parallel with a VIC-labelled reference gene assay (*Dot1l*, sourced from ThermoFisher) set at 2 copies (CNV2) on the Bio-Rad QX200 ddPCR system (Bio-Rad, CA) as per Codner and colleagues (Codner et al., 2016). Reaction mixes (22 μl) contained 2 μl crude DNA lysate or 50 ng of phenol/chloroform purified genomic DNA, 1x ddPCR Supermix for probes (Bio-Rad, CA, USA), 225 nM of each primer (two primers per assay) and 50 nM of each probe (one VIC-labelled probe for the reference gene assay and one FAM-labelled for the ssODN sequence assay). These reaction mixes were loaded either into DG8 cartridges together with 70 μl droplet oil per sample and droplets generated using the QX100 Droplet Generator or loaded in plate format into the Bio-Rad QX200 AutoDG and droplets generated as per the manufacturer’s instructions. Post droplet generation, the oil/reagent emulsion was transferred to a 96 well semi-skirted plate (Eppendorf AG, Hamburg, Germany) and the samples were amplified on the Bio-Rad C1000 Touch thermocycler (95°C for 10 min, followed by 40 cycles of 94°C for 30 s and 58°C for 60 s, with a final elongation step of 98 °C for 10 min, all temperature ramping set to 2.5°C/second). The plate containing the droplet amplicons was subsequently loaded into the QX200 Droplet Reader (Bio-Rad, CA, USA). Standard reagents and consumables supplied by Bio-Rad were used, including cartridges and gaskets, droplet generation oil and droplet reader oil. Copy number was assessed using the Quantasoft software using at least 10,000 accepted droplets per sample. Copy numbers were calculated by applying Poisson statistics to the fraction of end-point positive reactions and the 95% confidence interval of this measurement is shown.

## Abbreviations

bp: base-pair
Cas9: CRISPR associated protein 9
CRISPR: clustered regularly interspaced short palindromic repeat
ddPCR: droplet digital PCR
DNA: deoxyribonucleic acid
HA: homology arm
kb: kilobases
KI: knock-in
lssDNA: long single-stranded DNA
ONT: Oxford Nanopore Technologies
sgRNA: single guide RNA
SNP: single nucleotide polymorphism
ssODN: single-stranded oligo-deoxynucleotide
WT: wild type.

## Authors’ contributions

CM, GC, AA, AC, JL, MM, EM, JM, FP and LT designed protocols, sgRNA and oligonucleotides, synthesised reagents and genotyped progeny mice. CM performed MinION sequencing. CM, NDS and LT analysed long reads sequencing data. MH, MES and SW undertook mouse colony management. GC, HG, NDS and LT conceived the study. CM, GC, EM, MES and LT analysed the data. All authors contributed to the writing of the manuscript. All authors read and approved the final manuscript.

## Acknowledgements

The authors would like to thank the staff of the Mary Lyon Centre for providing excellent animal husbandry and microinjection services, Michael Micorescu (Oxford Nanopore Technologies, New York) and Duncan Sneddon for expert support with data mining, Dr Rosie Bunton-Stasyshyn for helpful discussions and Dr Louise Tinsley for expert assistance with the preparation of this manuscript.

## Ethics approval and consent to participate

All animal studies were licensed by the Home Office under the Animals (Scientific Procedures) Act 1986 Amendment Regulations 2012 (SI 4 2012/3039), UK, and additionally approved by the Institutional Ethical Review Committee.

## Availability of data and materials

ONT sequencing data deposited at the European Nucleotide Archive (https://www.ebi.ac.uk/ena) Identifiers are shown in Supplemental Table 1. All other data and their analyses are included in this published article and its supplementary information files.

## Supplemental Tables

*Supplemental Table ST1*: ONT sequencing experiments.

The table summarises the barcode, project, animal employed in each Nanopore sequencing experiment and the corresponding references to access datasets in the ENA repository.

*Supplemental Table ST2*: The table details the sequences of sgRNAs, for lssDNA templates, primers and probes employed in this study.

*Supplemental Table ST3*: The table details the mice that were obtained from each microinjection session during the course of this study.

*Supplemental Table ST4*: The table details the mouse colonies that were established during the course of this study.

## Supplemental Figures

*Supplemental Figure S1*: Experimental plan: Genome-edited loci can be characterised by Sanger sequencing (process highlighted in orange boxes), producing partial reads that must be assembled to reconstitute the whole region of interest. Alternatively, ONT sequencing (process highlighted in blue boxes) produces longer sequence reads spanning the whole region of interest.

*Supplemental Figure S2*: Required sequencing coverage for full identification of a known sequence. (a) shows the number of reads retained for by filters for each experiment. Note that the number of reads retained drops sharply for quality greater than q90. (b) shows the percentage of the sequenced interval identified by ONT sequencing for each WT segment analysed in Experiment A. Note that the large numbers of reads in lower quality range identify more reliably the reference sequences than fewer higher quality reads.

*Supplemental Figure S3*: Details of the analysis of the microinjection session containing animals interrogated by ONT sequencing for the Mpeg1 Cre project. The figure shows the PCR amplification of the genomic region of interest with (a) Mpeg1-F1 and Mpeg1-R1 (WT yields 873 bp amplicon, Cre KI yields 2412 bp amplicon) and (b) CreF and CreR primers (Cre KI yields 472 bp amplicon) from biopsies taken from the G_0_ animals. Animals yielding amplicons with CreF and CreR primers were subject to PCR to assess whether Cre is on target with primer combinations (c) Mpeg1-F1 and CreR (Cre KI yields 1440 bp amplicon) and (d) CreF and Mpeg1-R1 (Cre KI yields 1182 bp amplicon). (e) The panels show the sequencing of PCR amplicons using Mpeg1_F1/CreR and CreF/Mpeg1_R1 respectively obtained from animal Mpeg1-80. (f) The table details the analysis of G_0_ animals analysed: Animal ID, outcome of PCR analysis of the region of interest and the overall conclusion for each individual are shown. Panel of PCR amplicons with four different primer combinations (Mpeg1-F1 and Mpeg1-R1; CreF and CreR; Mpeg1-F1 and CreR; CreF and Mpeg1-R1) obtained for G_1_ animals derived from founder Mpeg1-75 crossed to WT (g) and founder Mpeg1-80 mated with WT (h). The table (i) details the ID, outcome of sequencing the region of interest, copy counting of the region of interest and the conclusion for each G_1_ individual. Sanger sequencing traces obtained from PCR amplification using Mpeg1_F1/CreR and CreF/Mpeg1_R1 respectively obtained from animal Mpeg1-75.1d. + is positive control amplified from an unrelated (a) WT, (b) Cre-KI animal. L1 = 1 kb DNA molecular weight ladder (thick band is 3 kb). Animal(s) interrogated by ONT sequence analysis are highlighted in green. * denotes animals with evidence of Cre KI on target but yielding multiple possible KI-related bands.

*Supplemental Figure S4*: Details of the analysis of the microinjection session and F1 colony containing animals interrogated by ONT sequencing for the Cx3cl1-flox project. The figure shows the PCR amplification of the genomic region of interest with (a) Cx3cl1-F1 and Cx3cl1-R1 primers (WT yields 1488 bp amplicon, floxed allele yields 1483 bp amplicon) and (b) LoxPF and LoxPR primers (floxed allele yields 835 bp amplicon) from biopsies taken from the G_0_ animals. (c) The panels show the sequencing of PCR amplicon obtained from animal Cx3cl1-10 with Cx3cl1-F1 and Cx3cl1-R1. LoxP site sequences are highlighted in blue. (d) The table details the G_0_ animals obtained from the microinjection. The ID and outcome of PCR analysis of the region of interest, as well as the conclusion for each founder are shown. Founder Cx3cl1-10 was mated for floxed allele transmission (LoxP PCR positive and sequence of complex mosaic). PCR amplification of region of interest with (e) Cx3cl1-F1 and Cx3cl1-R1 primers (1483 bp amplicon) and LoxPF and LoxPR primers (835 bp amplicon) from biopsies taken from Cx3cl1-10’s offspring. (f) The table details the first litter obtained by mating Cx3cl1-10 with a WT mouse. The ID, outcome of sequencing the region of interest and copy counting of the region of interest as well as the conclusion for each individual are shown. (g) Sanger sequence traces of the Cx3cl1 PCR product from G_1_ animal Cx3cl1-10.1c illustrating insertion of each LoxP site and associated genotyping handles (primer sequence and restriction enzyme site) highlighted in blue on target. + is positive control amplified from an unrelated (a) WT, (b) floxed animal. L1 = 1 kb DNA molecular weight ladder (thick band is 3 kb). Animal(s) interrogated by ONT sequence analysis are highlighted in green.

*Supplemental Figure S5*: General design strategy and generation of lssDNA donors for the generation of floxed alleles and cre KIs.

*Supplemental Figure S6*: Details of the analysis of the microinjection session containing animals interrogated by ONT sequencing for the Prdm8-flox project. The figure shows the PCR amplification of the genomic region of interest with (a) Prdm8-F1 and Prdm8-R1 primers (WT yields 1984 bp amplicon, floxed yields 2054 bp amplicon) and (b) LoxPF and LoxPR primers (floxed yields 1025 bp amplicon) from biopsies taken from the G_0_ animals. (c) The panels show the sequencing of PCR amplicon obtained from animal Prdm8-7 with Prdm8-F1 and LoxPF. LoxP site sequences are highlighted in blue. The SNP in the designed mutant sequence at 5’ end of the LoxPR primer sequence is highlightd in red. (d) The table details the G_0_ animals obtained from the microinjection analysed by ONT. The ID and outcome of PCR analysis of the region of interest, as well as the conclusion for each individual are shown. Animal(s) interrogated by ONT sequence analysis are highlighted in green. + is positive control amplified from an unrelated (a) WT, (b) floxed animal. L1 = 1 kb DNA molecular weight ladder (thick band is 3 kb), L2 = 100 bp DNA molecular weight ladder (thick bands are 1 kb and 500 bp).

*Supplemental Figure S7*: Details of the analysis of the microinjection session containing animals interrogated by ONT sequencing for the Pam-flox project. The figure shows the PCR amplification of the genomic region of interest with (a) Pam-F1 and Pam-R1 primers (WT yields 1426 bp amplicon, floxed yields 1431 b amplicon) and (b) LoxPF and LoxPR primers (floxed yields 801 bp amplicon) from biopsies taken from the G_0_ animals. (c) The panels show the sequencing of PCR amplicon obtained from animal Pam-3 with Pam-F1 and Pam-R1. LoxP site sequences are highlighted in blue. (d) The table details the G_0_ animals analysed: ID, outcome of PCR analysis of the region of interest and conclusion for each individual are shown. PCR amplification of region of interest with (e) Pam-F1 and Pam-R1 primers (1431 bp amplicon) and (f) LoxPF and LoxPR primers (801 bp amplicon) from biopsies taken from Pam-3’s offspring. (g) The table details the first litter obtained by mating Pam-flox-3 with a WT mouse. The ID, outcome of PCR amplification of the regions of interest as well as the initial conclusion for each individual are shown. NB. No Sanger sequencing was performed on the Pam floxed G_1_ generation prior to analysis with ONT. Animal(s) interrogated by ONT sequence analysis are highlighted in green. + is positive control amplified from an unrelated (a) WT, (b) floxed animal. L1 = 1 kb DNA molecular weight ladder (thick band is 3 kb).

*Supplemental Figure S8*: Details of the analysis of the microinjection session containing animals interrogated by ONT sequencing for the Hnf1a-flox project. The figure shows the PCR amplification of the genomic region of interest with (a) Hnf1a-F1 and Hnf1a-R1 primers (WT yields 1221 bp amplicon, floxed allele yields 1191 bp amplicon) and (b) LoxPF and LoxPR primers (floxed allele yields 691 bp amplicon) from biopsies taken from the G_0_ animals. (c) The panels show the sequencing of PCR amplicon obtained from animal Hnf1a-66 with Hnf1a-F1 and Hnf1a-R1 and sequenced with LoxPF and LoxPR. LoxP site sequences are highlighted in blue. (d) The table details the G_0_ animals obtained; the ID and outcome of PCR analysis of the region of interest, as well as the conclusion for each individual are shown. Animal(s) interrogated by ONT sequence analysis are highlighted in green. + is positive control amplified from an unrelated (a) WT, (b) floxed animal. L1 = 1 kb DNA molecular weight ladder (thick band is 3 kb), L2 = 100 bp DNA molecular weight ladder (thick bands are 1 kb and 500 bp).

*Supplemental Figure S9*: Details of the analysis of the microinjection sessions containing animals interrogated by ONT sequencing for the Inpp5k-flox project. The figure shows the PCR amplification of the genomic region of interest with (a) Inpp5k-F1 and Inpp5k-R1 primers (WT yields 1701 bp amplicon, floxed allele yields 1705 bp amplicon) and (b) LoxPF and LoxPR primers (floxed allele yields 1194 bp amplicon) from biopsies taken from the G_0_ animals. (c) The panels show the sequencing of PCR amplicon obtained from animal Inpp5k-7 with Inpp5k-F1 and LoxPR, and with LoxPF and Inpp5k-R1 respectively. LoxP site sequences are highlighted in blue. (d) The table details the G_0_ animals analysed: the ID, outcome of PCR analysis of the region of interest and the conclusion for each individual are shown. Animal(s) interrogated by ONT sequence analysis are highlighted in green. + is positive control amplified from an unrelated (a) WT, (b) floxed animal. L1 = 1 kb DNA molecular weight ladder (thick band is 3 kb).

*Supplemental Figure S10*: Confirmation of potential animals for the Pam-flox and Hnf1a-floxed projects. (a) shows the outcome of ONT sequencing of the founder animal Pam-flox-3 aligned against the mutant Pam-flox reference visualised with IGV. (b) shows the outcome of ONT sequencing of the founder animal Hnf1a-flox-66 aligned against the mutant Hnf1a-flox reference visualised with IGV. The alignments reflects the noisy nature of the method with errors distributed across the length of the sequenced segment. Note the complete alignment of reads to designed mutant sequence (grey histograms).

*Supplemental Figure S11:* Alignment of sequencing reads from G_0_s Inpp5k-7 and -33 against the Inpp5k-flox reference without filtering for determinants. The yellow frame highlights the presence of segments that are different to the reference in G_0_ Inpp5k-7. The blue and red frames highlight that although G_0_ Inpp5k-36 contains sequences that are overall similar to the mutant sequence reference, these alleles also contain point mutations.

*Supplemental Figure S12*: Details of the analysis of the G_1_ generation containing animals interrogated by ONT sequencing for the Inpp5k-flox project. The figure shows the PCR amplification of the genomic region of interest with (a) Inpp5k-F1 and Inpp5k-R1 primers (WT yields 1701 bp amplicon, floxed allele yields 1705 bp amplicon) and (b) LoxPF and LoxPR primers (floxed allele yields 1194 bp amplicon) from biopsies taken from the G_1_ animals derived from crossing founder animals Inpp5k-7 and Inpp5k-8 to WT. (c) The table details the G_1_ animals obtained from the two lines. The ID and outcome of PCR analysis of the region of interest, as well as the conclusion for each individual are shown. (d) The panels show the sequencing of PCR amplicon obtained from animal Inpp5k-7.1b with Inpp5k-F1 and LoxPR, and with LoxPF and Inpp5k-R1 respectively. Deviations from the intended mutant sequence are highlighted in blue. Animal(s) interrogated by ONT sequence analysis are highlighted in green. + is positive control amplified from an unrelated (a) WT, (b) floxed animal. L1 = 1 kb DNA molecular weight ladder (thick band is 3 kb).

*Supplemental Figure S13:* Details of the analysis microinjection session containing animals interrogated by ONT sequencing for the 6430573F11Rik project. The figure shows the PCR amplification of the genomic region of interest with (a) 6430573F11Rik-F3 and 6430573F11Rik-R2 primers (WT yields 1724 bp amplicon, floxed yields 1721 bp amplicon) and (b) LoxPF and LoxPR primers (floxed yields 999 bp amplicon) from biopsies taken from the G0 animals. (c) The panels show the sequencing of PCR amplicon obtained from animal 6430573F11Rik-11 with 6430573F11Rik-F2 and 6430573F11Rik-R3. LoxP site sequences are highlighted in blue. The SNP in the critical region is also highlighted in blue (d) The table details the G0 animals analysed: The ID, outcome of PCR analysis of the region of interest and the conclusion for each individual are shown. Animal(s) interrogated by ONT sequence analysis are highlighted in green. + is positive control amplified from an unrelated (a) WT, (b) floxed animal. L1 = 1 kb DNA molecular weight ladder (thick band is 3 kb).

