## Supplemental Figures S1 to S13 for "Application of long-read sequencing for robust identification of correct alleles in genome edited animals"

### Slide 1
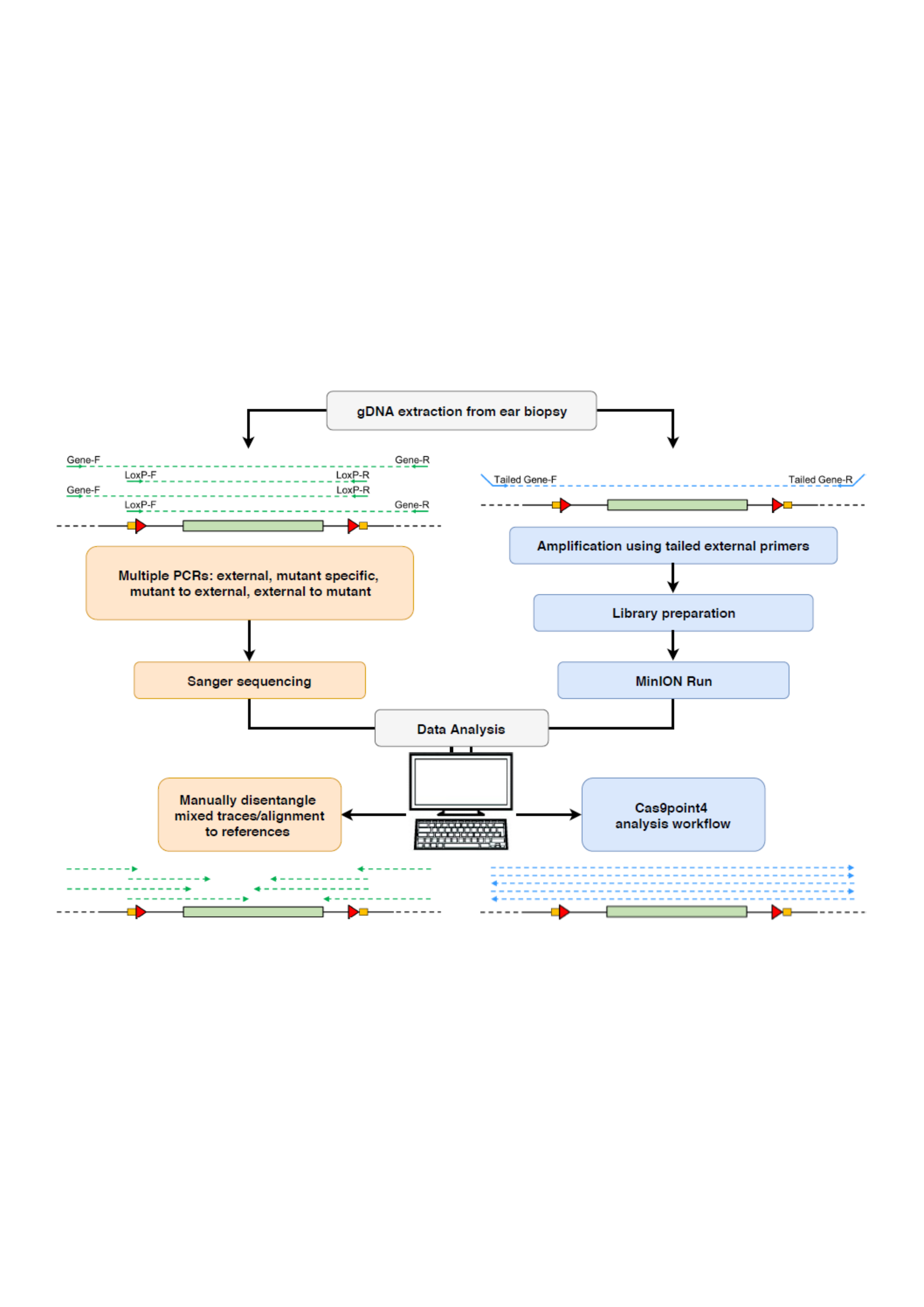

### Slide 2
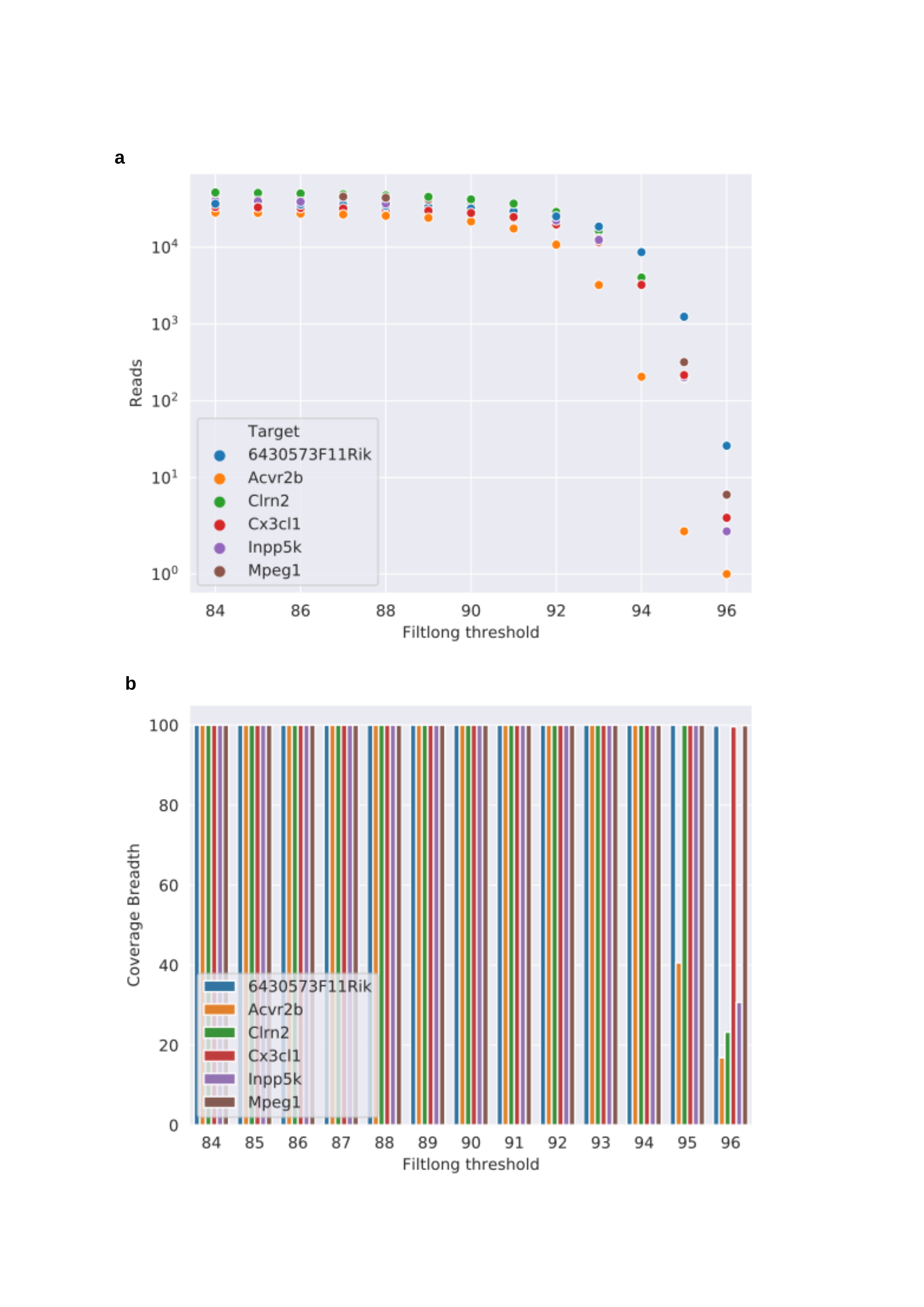

a
b

### Slide 3
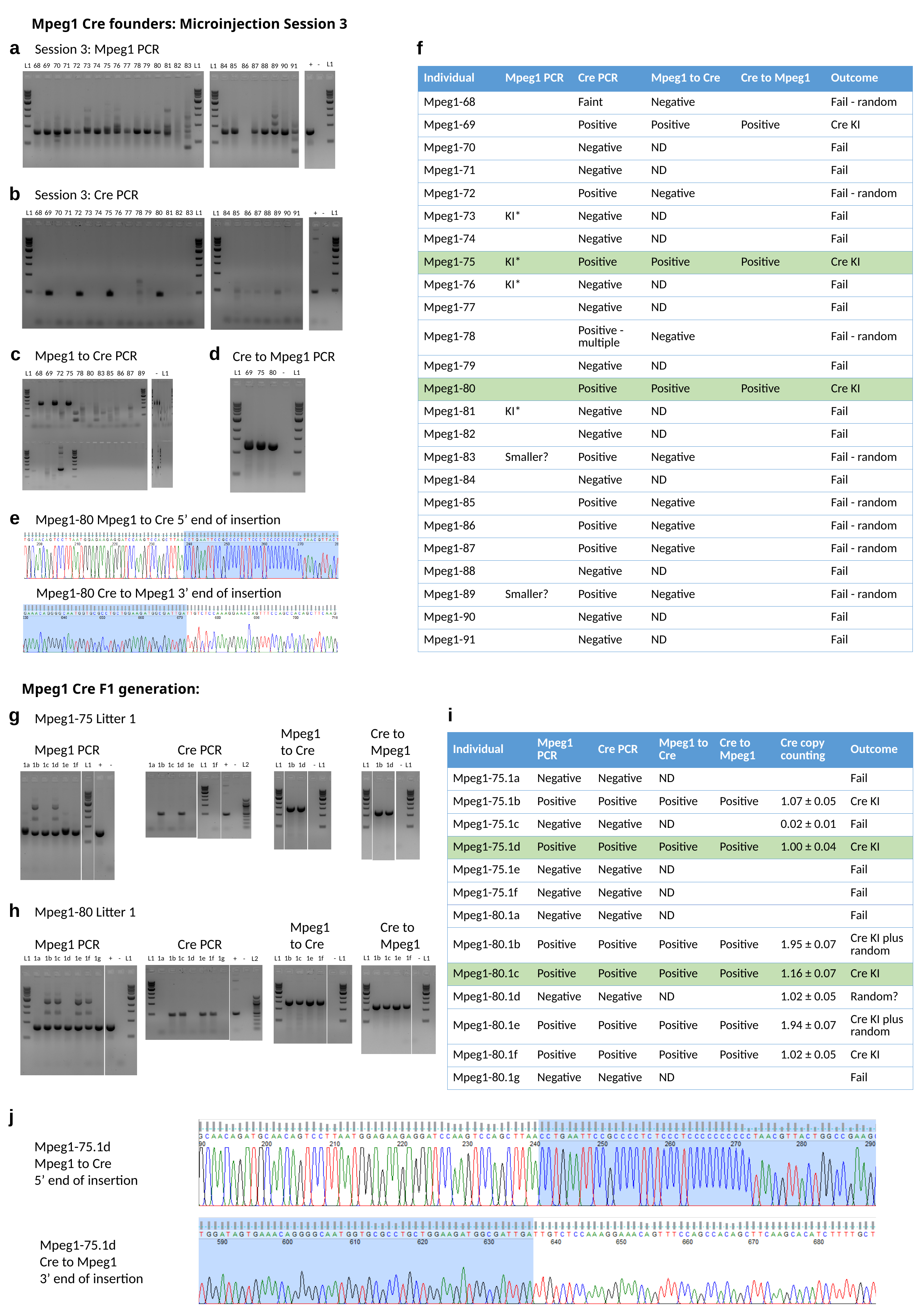

Mpeg1 Cre founders: Microinjection Session 3
a
f
Session 3: Mpeg1 PCR
L1
+
-
L1
68
69
70
71
72
73
74
75
76
77
78
79
80
81
82
83
L1
L1
84
85
86
87
88
89
90
91
| Individual | Mpeg1 PCR | Cre PCR | Mpeg1 to Cre | Cre to Mpeg1 | Outcome |
| --- | --- | --- | --- | --- | --- |
| Mpeg1-68 | | Faint | Negative | | Fail - random |
| Mpeg1-69 | | Positive | Positive | Positive | Cre KI |
| Mpeg1-70 | | Negative | ND | | Fail |
| Mpeg1-71 | | Negative | ND | | Fail |
| Mpeg1-72 | | Positive | Negative | | Fail - random |
| Mpeg1-73 | KI\* | Negative | ND | | Fail |
| Mpeg1-74 | | Negative | ND | | Fail |
| Mpeg1-75 | KI\* | Positive | Positive | Positive | Cre KI |
| Mpeg1-76 | KI\* | Negative | ND | | Fail |
| Mpeg1-77 | | Negative | ND | | Fail |
| Mpeg1-78 | | Positive - multiple | Negative | | Fail - random |
| Mpeg1-79 | | Negative | ND | | Fail |
| Mpeg1-80 | | Positive | Positive | Positive | Cre KI |
| Mpeg1-81 | KI\* | Negative | ND | | Fail |
| Mpeg1-82 | | Negative | ND | | Fail |
| Mpeg1-83 | Smaller? | Positive | Negative | | Fail - random |
| Mpeg1-84 | | Negative | ND | | Fail |
| Mpeg1-85 | | Positive | Negative | | Fail - random |
| Mpeg1-86 | | Positive | Negative | | Fail - random |
| Mpeg1-87 | | Positive | Negative | | Fail - random |
| Mpeg1-88 | | Negative | ND | | Fail |
| Mpeg1-89 | Smaller? | Positive | Negative | | Fail - random |
| Mpeg1-90 | | Negative | ND | | Fail |
| Mpeg1-91 | | Negative | ND | | Fail |
b
Session 3: Cre PCR
L1
+
-
L1
68
69
70
71
72
73
74
75
76
77
78
79
80
81
82
83
L1
L1
84
85
86
87
88
89
90
91
d
c
Mpeg1 to Cre PCR
Cre to Mpeg1 PCR
L1
69
75
80
-
L1
L1
68
69
72
75
78
80
83
85
86
87
89
-
L1
e
Mpeg1-80 Mpeg1 to Cre 5’ end of insertion
Mpeg1-80 Cre to Mpeg1 3’ end of insertion
Mpeg1 Cre F1 generation:
g
i
Mpeg1-75 Litter 1
Mpeg1
to Cre
Cre to
Mpeg1
| Individual | Mpeg1 PCR | Cre PCR | Mpeg1 to Cre | Cre to Mpeg1 | Cre copy counting | Outcome |
| --- | --- | --- | --- | --- | --- | --- |
| Mpeg1-75.1a | Negative | Negative | ND | | | Fail |
| Mpeg1-75.1b | Positive | Positive | Positive | Positive | 1.07 ± 0.05 | Cre KI |
| Mpeg1-75.1c | Negative | Negative | ND | | 0.02 ± 0.01 | Fail |
| Mpeg1-75.1d | Positive | Positive | Positive | Positive | 1.00 ± 0.04 | Cre KI |
| Mpeg1-75.1e | Negative | Negative | ND | | | Fail |
| Mpeg1-75.1f | Negative | Negative | ND | | | Fail |
| Mpeg1-80.1a | Negative | Negative | ND | | | Fail |
| Mpeg1-80.1b | Positive | Positive | Positive | Positive | 1.95 ± 0.07 | Cre KI plus random |
| Mpeg1-80.1c | Positive | Positive | Positive | Positive | 1.16 ± 0.07 | Cre KI |
| Mpeg1-80.1d | Negative | Negative | ND | | 1.02 ± 0.05 | Random? |
| Mpeg1-80.1e | Positive | Positive | Positive | Positive | 1.94 ± 0.07 | Cre KI plus random |
| Mpeg1-80.1f | Positive | Positive | Positive | Positive | 1.02 ± 0.05 | Cre KI |
| Mpeg1-80.1g | Negative | Negative | ND | | | Fail |
Mpeg1 PCR
Cre PCR
+ - L2
1a
1b
1c
1d
1e
1f
L1
+ -
1a
1b
1c
1d
1e
1f
1b
1d
1b
1d
L1
L1
- L1
L1
- L1
h
Mpeg1-80 Litter 1
Mpeg1
to Cre
Cre to
Mpeg1
Mpeg1 PCR
Cre PCR
1b
1c
1e
1f
L1
L1
1a
1b
1c
1d
1e
1f
1g
+ - L1
L1
1a
1b
1c
1d
1e
1f
1g
+ - L2
1b
1c
1e
1f
L1
- L1
- L1
j
Mpeg1-75.1d
Mpeg1 to Cre
5’ end of insertion
Mpeg1-75.1d
Cre to Mpeg1
3’ end of insertion

### Slide 4
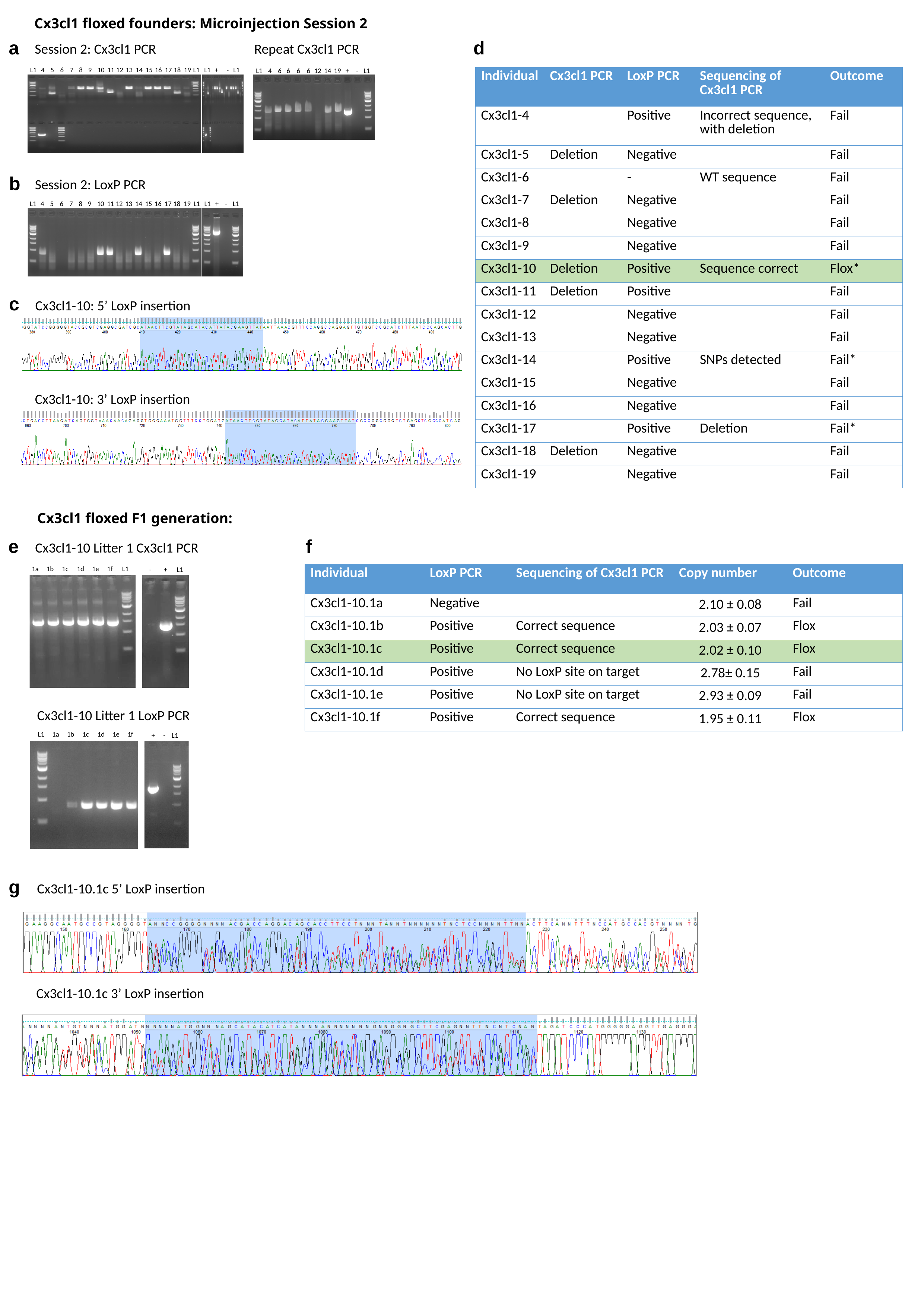

Cx3cl1 floxed founders: Microinjection Session 2
a
d
Session 2: Cx3cl1 PCR
Repeat Cx3cl1 PCR
L1
4
5
6
7
8
9
10
11
12
13
14
15
16
17
18
19
L1
L1
+
-
L1
L1
4
6
6
6
6
12
14
19
+
-
L1
| Individual | Cx3cl1 PCR | LoxP PCR | Sequencing of Cx3cl1 PCR | Outcome |
| --- | --- | --- | --- | --- |
| Cx3cl1-4 | | Positive | Incorrect sequence, with deletion | Fail |
| Cx3cl1-5 | Deletion | Negative | | Fail |
| Cx3cl1-6 | | - | WT sequence | Fail |
| Cx3cl1-7 | Deletion | Negative | | Fail |
| Cx3cl1-8 | | Negative | | Fail |
| Cx3cl1-9 | | Negative | | Fail |
| Cx3cl1-10 | Deletion | Positive | Sequence correct | Flox\* |
| Cx3cl1-11 | Deletion | Positive | | Fail |
| Cx3cl1-12 | | Negative | | Fail |
| Cx3cl1-13 | | Negative | | Fail |
| Cx3cl1-14 | | Positive | SNPs detected | Fail\* |
| Cx3cl1-15 | | Negative | | Fail |
| Cx3cl1-16 | | Negative | | Fail |
| Cx3cl1-17 | | Positive | Deletion | Fail\* |
| Cx3cl1-18 | Deletion | Negative | | Fail |
| Cx3cl1-19 | | Negative | | Fail |
b
Session 2: LoxP PCR
L1
4
5
6
7
8
9
10
11
12
13
14
15
16
17
18
19
L1
L1
+
-
L1
c
Cx3cl1-10: 5’ LoxP insertion
Cx3cl1-10: 3’ LoxP insertion
Cx3cl1 floxed F1 generation:
e
f
Cx3cl1-10 Litter 1 Cx3cl1 PCR
1a
1b
1c
1d
1e
1f L1
-
+ L1
| Individual | LoxP PCR | Sequencing of Cx3cl1 PCR | Copy number | Outcome |
| --- | --- | --- | --- | --- |
| Cx3cl1-10.1a | Negative | | 2.10 ± 0.08 | Fail |
| Cx3cl1-10.1b | Positive | Correct sequence | 2.03 ± 0.07 | Flox |
| Cx3cl1-10.1c | Positive | Correct sequence | 2.02 ± 0.10 | Flox |
| Cx3cl1-10.1d | Positive | No LoxP site on target | 2.78± 0.15 | Fail |
| Cx3cl1-10.1e | Positive | No LoxP site on target | 2.93 ± 0.09 | Fail |
| Cx3cl1-10.1f | Positive | Correct sequence | 1.95 ± 0.11 | Flox |
Cx3cl1-10 Litter 1 LoxP PCR
L1 1a
1b
1c
1d
1e
1f
+
- L1
g
Cx3cl1-10.1c 5’ LoxP insertion
Cx3cl1-10.1c 3’ LoxP insertion

### Slide 5
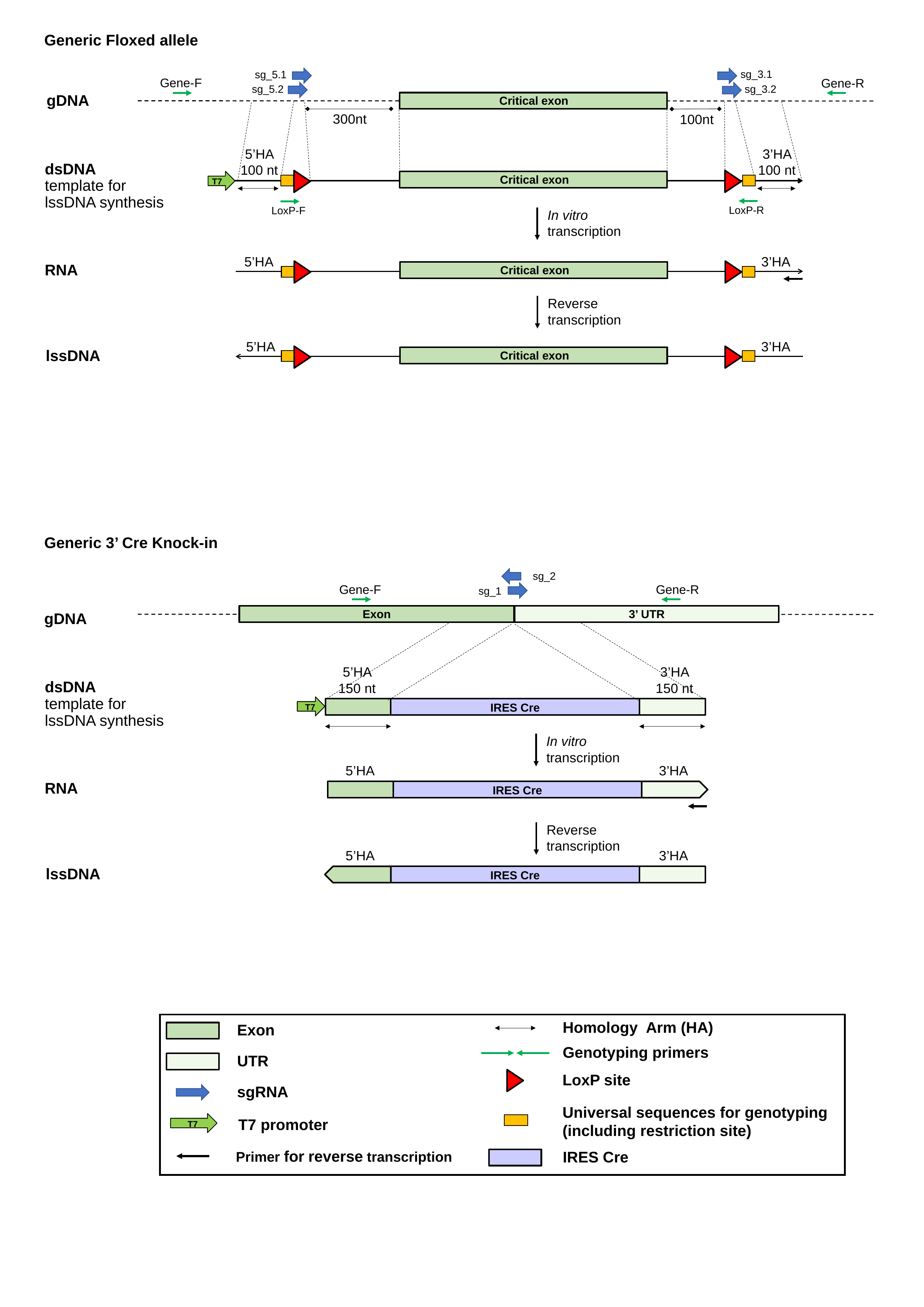

Generic Floxed allele
sg_3.1
sg_5.1
Gene-F
Gene-R
sg_5.2
sg_3.2
Critical exon
#### gDNA
300nt
100nt
3’HA 100 nt
5’HA
100 nt
T7
Critical exon
dsDNA
template for lssDNA synthesis
LoxP-R
LoxP-F
In vitro transcription
5’HA
3’HA
Critical exon
RNA
Reverse transcription
5’HA
3’HA
Critical exon
lssDNA
Generic 3’ Cre Knock-in
sg_2
Gene-F
Gene-R
sg_1
Exon
3’ UTR
gDNA
3’HA 150 nt
5’HA
150 nt
dsDNA
template for lssDNA synthesis
T7
IRES Cre
In vitro transcription
5’HA
3’HA
RNA
IRES Cre
Reverse transcription
5’HA
3’HA
IRES Cre
lssDNA
Homology Arm (HA)
Exon
Genotyping primers
UTR
LoxP site
sgRNA
Universal sequences for genotyping (including restriction site)
T7
T7 promoter
Primer for reverse transcription
IRES Cre

### Slide 6
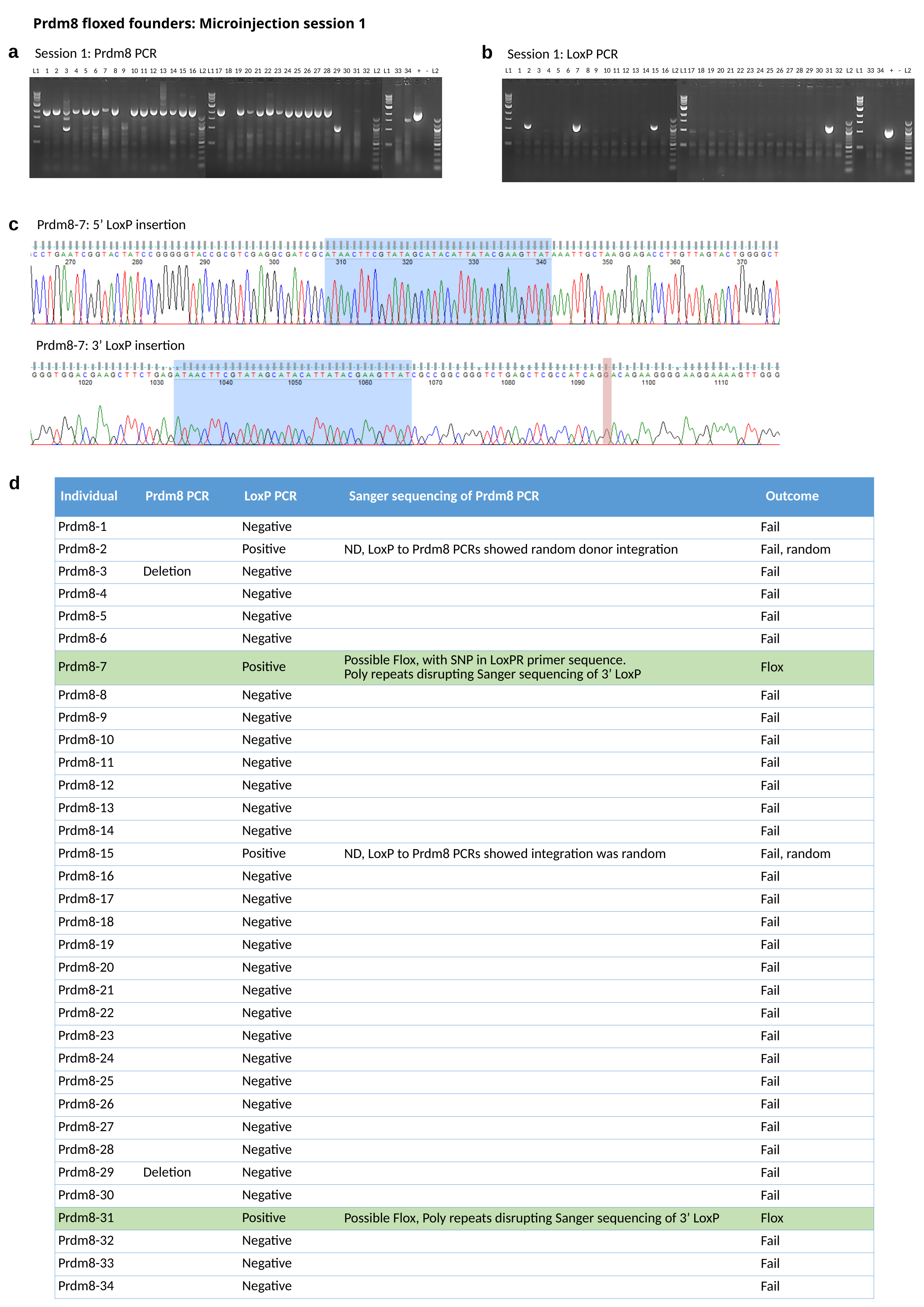

Prdm8 floxed founders: Microinjection session 1
a
b
Session 1: Prdm8 PCR
Session 1: LoxP PCR
L1
1
2
3
4
5
6
7
8
9
10
11
12
13
14
15
16
L2
L1
17
18
19
20
21
22
23
24
25
26
27
28
29
30
31
32
L2
L1
33
34
+
-
L2
L1
1
2
3
4
5
6
7
8
9
10
11
12
13
14
15
16
L2
L1
17
18
19
20
21
22
23
24
25
26
27
28
29
30
31
32
L2
L1
33
34
+
-
L2
c
Prdm8-7: 5’ LoxP insertion
Prdm8-7: 3’ LoxP insertion
d
| Individual | Prdm8 PCR | LoxP PCR | Sanger sequencing of Prdm8 PCR | Outcome |
| --- | --- | --- | --- | --- |
| Prdm8-1 | | Negative | | Fail |
| Prdm8-2 | | Positive | ND, LoxP to Prdm8 PCRs showed random donor integration | Fail, random |
| Prdm8-3 | Deletion | Negative | | Fail |
| Prdm8-4 | | Negative | | Fail |
| Prdm8-5 | | Negative | | Fail |
| Prdm8-6 | | Negative | | Fail |
| Prdm8-7 | | Positive | Possible Flox, with SNP in LoxPR primer sequence. Poly repeats disrupting Sanger sequencing of 3’ LoxP | Flox |
| Prdm8-8 | | Negative | | Fail |
| Prdm8-9 | | Negative | | Fail |
| Prdm8-10 | | Negative | | Fail |
| Prdm8-11 | | Negative | | Fail |
| Prdm8-12 | | Negative | | Fail |
| Prdm8-13 | | Negative | | Fail |
| Prdm8-14 | | Negative | | Fail |
| Prdm8-15 | | Positive | ND, LoxP to Prdm8 PCRs showed integration was random | Fail, random |
| Prdm8-16 | | Negative | | Fail |
| Prdm8-17 | | Negative | | Fail |
| Prdm8-18 | | Negative | | Fail |
| Prdm8-19 | | Negative | | Fail |
| Prdm8-20 | | Negative | | Fail |
| Prdm8-21 | | Negative | | Fail |
| Prdm8-22 | | Negative | | Fail |
| Prdm8-23 | | Negative | | Fail |
| Prdm8-24 | | Negative | | Fail |
| Prdm8-25 | | Negative | | Fail |
| Prdm8-26 | | Negative | | Fail |
| Prdm8-27 | | Negative | | Fail |
| Prdm8-28 | | Negative | | Fail |
| Prdm8-29 | Deletion | Negative | | Fail |
| Prdm8-30 | | Negative | | Fail |
| Prdm8-31 | | Positive | Possible Flox, Poly repeats disrupting Sanger sequencing of 3’ LoxP | Flox |
| Prdm8-32 | | Negative | | Fail |
| Prdm8-33 | | Negative | | Fail |
| Prdm8-34 | | Negative | | Fail |

### Slide 7
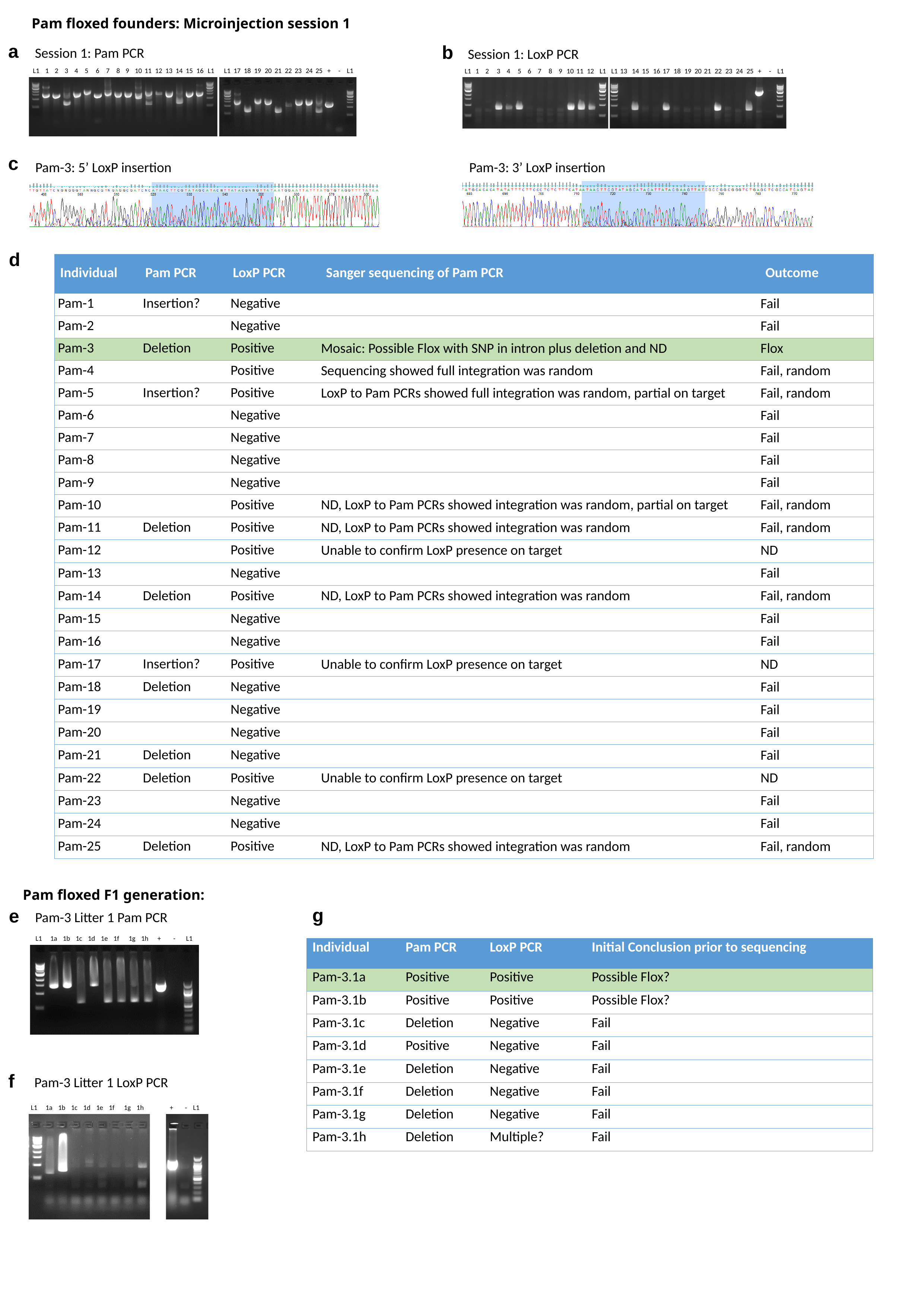

Pam floxed founders: Microinjection session 1
a
b
Session 1: Pam PCR
Session 1: LoxP PCR
L1
1
2
3
4
5
6
7
8
9
10
11
12
13
14
15
16
L1
L1
17
18
19
20
21
22
23
24
25
+
-
L1
L1
1
2
3
4
5
6
7
8
9
10
11
12
L1
L1
13
14
15
16
17
18
19
20
21
22
23
24
25
+
-
L1
c
Pam-3: 5’ LoxP insertion
Pam-3: 3’ LoxP insertion
d
| Individual | Pam PCR | LoxP PCR | Sanger sequencing of Pam PCR | Outcome |
| --- | --- | --- | --- | --- |
| Pam-1 | Insertion? | Negative | | Fail |
| Pam-2 | | Negative | | Fail |
| Pam-3 | Deletion | Positive | Mosaic: Possible Flox with SNP in intron plus deletion and ND | Flox |
| Pam-4 | | Positive | Sequencing showed full integration was random | Fail, random |
| Pam-5 | Insertion? | Positive | LoxP to Pam PCRs showed full integration was random, partial on target | Fail, random |
| Pam-6 | | Negative | | Fail |
| Pam-7 | | Negative | | Fail |
| Pam-8 | | Negative | | Fail |
| Pam-9 | | Negative | | Fail |
| Pam-10 | | Positive | ND, LoxP to Pam PCRs showed integration was random, partial on target | Fail, random |
| Pam-11 | Deletion | Positive | ND, LoxP to Pam PCRs showed integration was random | Fail, random |
| Pam-12 | | Positive | Unable to confirm LoxP presence on target | ND |
| Pam-13 | | Negative | | Fail |
| Pam-14 | Deletion | Positive | ND, LoxP to Pam PCRs showed integration was random | Fail, random |
| Pam-15 | | Negative | | Fail |
| Pam-16 | | Negative | | Fail |
| Pam-17 | Insertion? | Positive | Unable to confirm LoxP presence on target | ND |
| Pam-18 | Deletion | Negative | | Fail |
| Pam-19 | | Negative | | Fail |
| Pam-20 | | Negative | | Fail |
| Pam-21 | Deletion | Negative | | Fail |
| Pam-22 | Deletion | Positive | Unable to confirm LoxP presence on target | ND |
| Pam-23 | | Negative | | Fail |
| Pam-24 | | Negative | | Fail |
| Pam-25 | Deletion | Positive | ND, LoxP to Pam PCRs showed integration was random | Fail, random |
Pam floxed F1 generation:
g
e
Pam-3 Litter 1 Pam PCR
L1
1a
1b
1c
1d
1e
1f 1g
1h
+
-
L1
| Individual | Pam PCR | LoxP PCR | Initial Conclusion prior to sequencing |
| --- | --- | --- | --- |
| Pam-3.1a | Positive | Positive | Possible Flox? |
| Pam-3.1b | Positive | Positive | Possible Flox? |
| Pam-3.1c | Deletion | Negative | Fail |
| Pam-3.1d | Positive | Negative | Fail |
| Pam-3.1e | Deletion | Negative | Fail |
| Pam-3.1f | Deletion | Negative | Fail |
| Pam-3.1g | Deletion | Negative | Fail |
| Pam-3.1h | Deletion | Multiple? | Fail |
f
Pam-3 Litter 1 LoxP PCR
L1
1a
1b
1c
1d
1e
1f 1g
1h
+
-
L1

### Slide 8
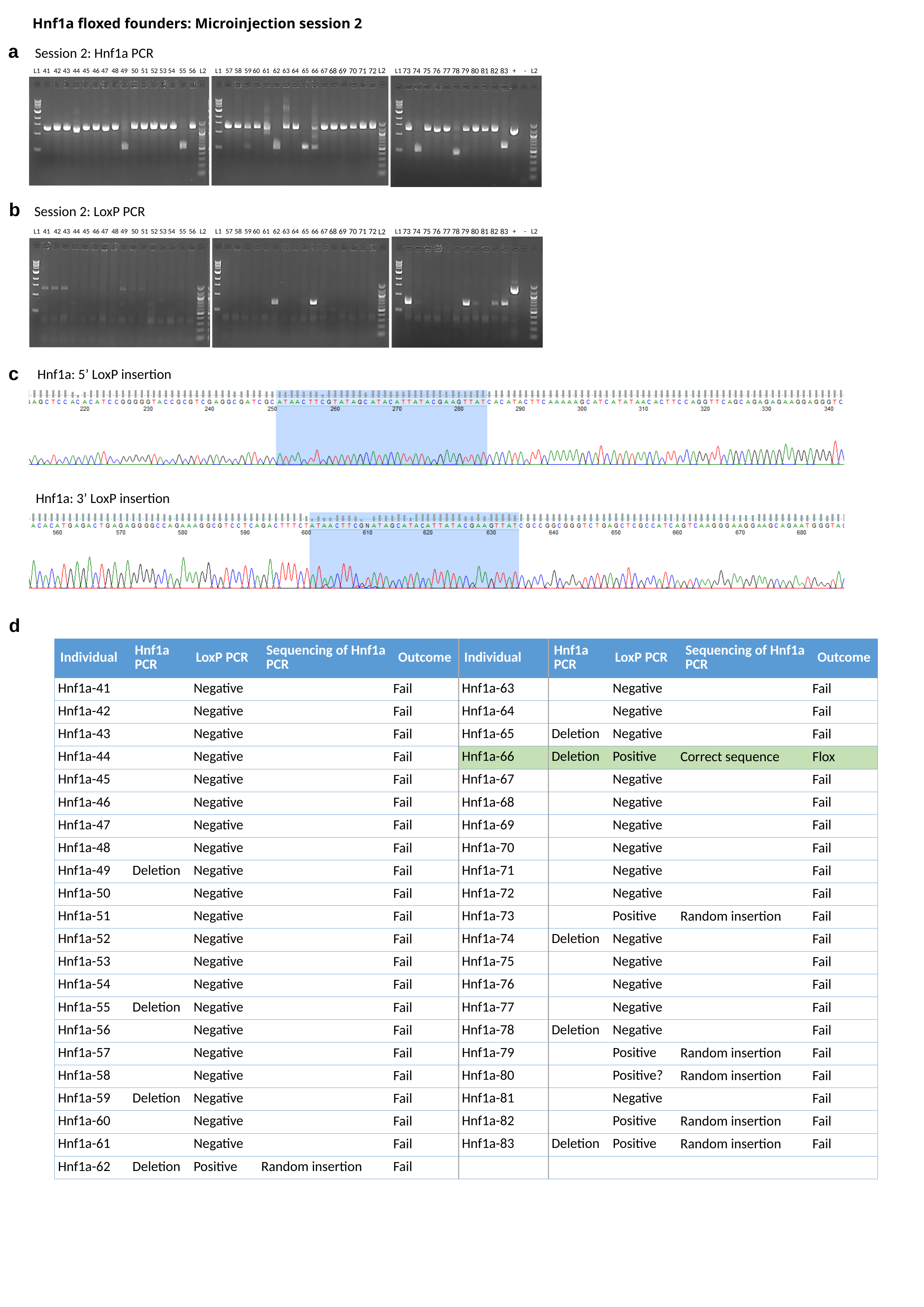

Hnf1a floxed founders: Microinjection session 2
a
Session 2: Hnf1a PCR
L2
68
69
70
71
72
73
74
75
76
77
78
79
80
81
82
83
L1
41
42
43
44
45
46
47
48
49
50
51
52
53
54
55
56
L2
L1
57
58
59
60
61
62
63
64
65
66
67
L1
+
-
L2
b
Session 2: LoxP PCR
68
69
70
71
72
73
74
75
76
77
78
79
80
81
82
83
L2
L1
41
42
43
44
45
46
47
48
49
50
51
52
53
54
55
56
L2
L1
57
58
59
60
61
62
63
64
65
66
67
L1
+
-
L2
c
Hnf1a: 5’ LoxP insertion
Hnf1a: 3’ LoxP insertion
d
| Individual | Hnf1a PCR | LoxP PCR | Sequencing of Hnf1a PCR | Outcome | Individual | Hnf1a PCR | LoxP PCR | Sequencing of Hnf1a PCR | Outcome |
| --- | --- | --- | --- | --- | --- | --- | --- | --- | --- |
| Hnf1a-41 | | Negative | | Fail | Hnf1a-63 | | Negative | | Fail |
| Hnf1a-42 | | Negative | | Fail | Hnf1a-64 | | Negative | | Fail |
| Hnf1a-43 | | Negative | | Fail | Hnf1a-65 | Deletion | Negative | | Fail |
| Hnf1a-44 | | Negative | | Fail | Hnf1a-66 | Deletion | Positive | Correct sequence | Flox |
| Hnf1a-45 | | Negative | | Fail | Hnf1a-67 | | Negative | | Fail |
| Hnf1a-46 | | Negative | | Fail | Hnf1a-68 | | Negative | | Fail |
| Hnf1a-47 | | Negative | | Fail | Hnf1a-69 | | Negative | | Fail |
| Hnf1a-48 | | Negative | | Fail | Hnf1a-70 | | Negative | | Fail |
| Hnf1a-49 | Deletion | Negative | | Fail | Hnf1a-71 | | Negative | | Fail |
| Hnf1a-50 | | Negative | | Fail | Hnf1a-72 | | Negative | | Fail |
| Hnf1a-51 | | Negative | | Fail | Hnf1a-73 | | Positive | Random insertion | Fail |
| Hnf1a-52 | | Negative | | Fail | Hnf1a-74 | Deletion | Negative | | Fail |
| Hnf1a-53 | | Negative | | Fail | Hnf1a-75 | | Negative | | Fail |
| Hnf1a-54 | | Negative | | Fail | Hnf1a-76 | | Negative | | Fail |
| Hnf1a-55 | Deletion | Negative | | Fail | Hnf1a-77 | | Negative | | Fail |
| Hnf1a-56 | | Negative | | Fail | Hnf1a-78 | Deletion | Negative | | Fail |
| Hnf1a-57 | | Negative | | Fail | Hnf1a-79 | | Positive | Random insertion | Fail |
| Hnf1a-58 | | Negative | | Fail | Hnf1a-80 | | Positive? | Random insertion | Fail |
| Hnf1a-59 | Deletion | Negative | | Fail | Hnf1a-81 | | Negative | | Fail |
| Hnf1a-60 | | Negative | | Fail | Hnf1a-82 | | Positive | Random insertion | Fail |
| Hnf1a-61 | | Negative | | Fail | Hnf1a-83 | Deletion | Positive | Random insertion | Fail |
| Hnf1a-62 | Deletion | Positive | Random insertion | Fail | | | | | |

### Slide 9
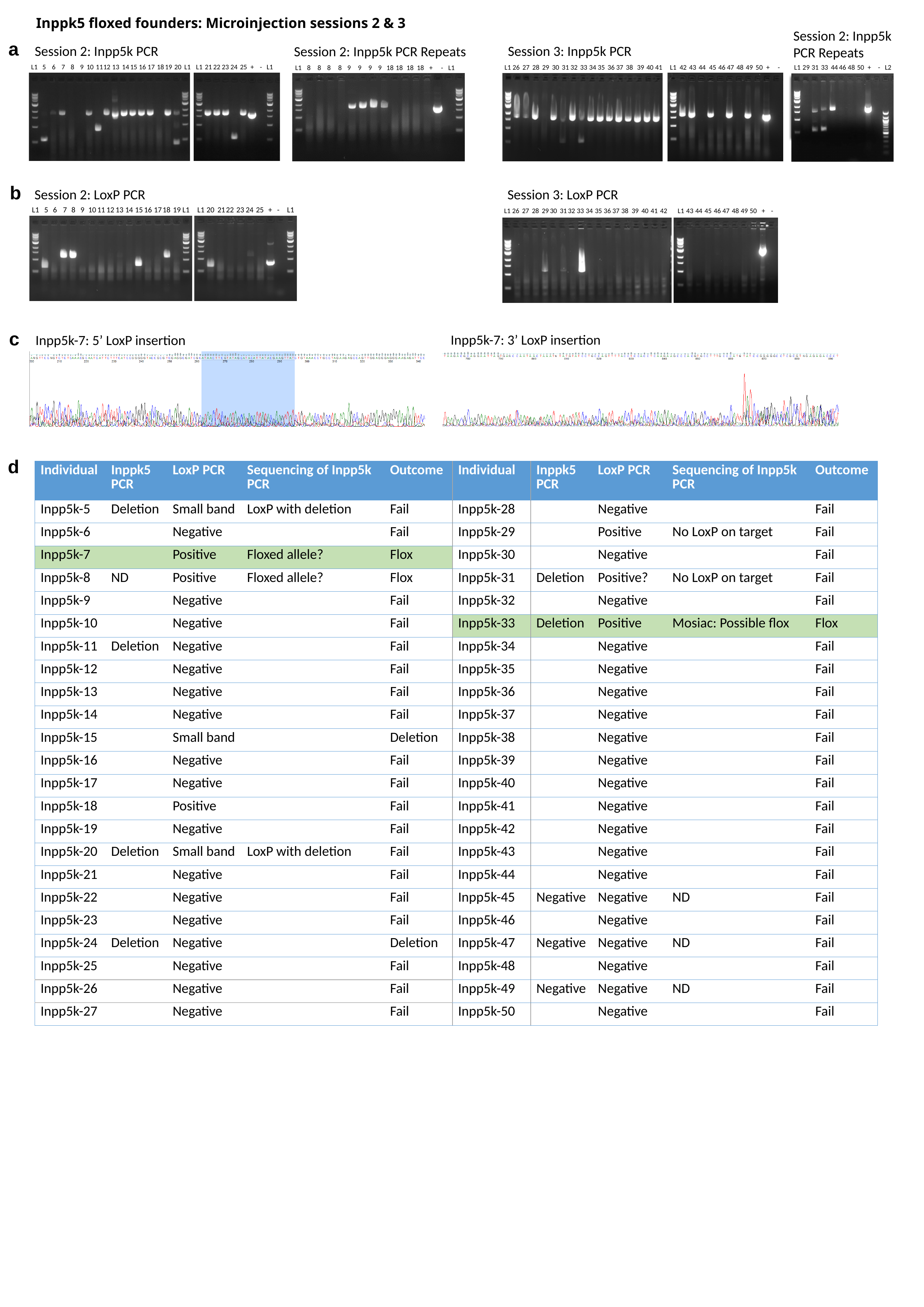

Inppk5 floxed founders: Microinjection sessions 2 & 3
Session 2: Inpp5k PCR Repeats
a
Session 3: Inpp5k PCR
Session 2: Inpp5k PCR
Session 2: Inpp5k PCR Repeats
L1
5
6
7
8
9
10
11
12
13
14
15
16
17
18
19
20
L1
L1
21
22
23
24
25
+
-
L1
L1
26
27
28
29
30
31
32
33
34
35
36
37
38
39
40
41
L1
42
43
44
45
46
47
48
49
50
+
-
L1
29
31
33
44
46
48
50
+
-
L2
L1
8
8
8
8
9
9
9
9
18
18
18
18
+
-
L1
b
Session 2: LoxP PCR
Session 3: LoxP PCR
L1
5
6
7
8
9
10
11
12
13
14
15
16
17
18
19
L1
L1
20
21
22
23
24
25
+
-
L1
L1
26
27
28
29
30
31
32
33
34
35
36
37
38
39
40
41
42
L1
43
44
45
46
47
48
49
50
+
-
c
Inpp5k-7: 3’ LoxP insertion
Inpp5k-7: 5’ LoxP insertion
d
| Individual | Inppk5 PCR | LoxP PCR | Sequencing of Inpp5k PCR | Outcome | Individual | Inppk5 PCR | LoxP PCR | Sequencing of Inpp5k PCR | Outcome |
| --- | --- | --- | --- | --- | --- | --- | --- | --- | --- |
| Inpp5k-5 | Deletion | Small band | LoxP with deletion | Fail | Inpp5k-28 | | Negative | | Fail |
| Inpp5k-6 | | Negative | | Fail | Inpp5k-29 | | Positive | No LoxP on target | Fail |
| Inpp5k-7 | | Positive | Floxed allele? | Flox | Inpp5k-30 | | Negative | | Fail |
| Inpp5k-8 | ND | Positive | Floxed allele? | Flox | Inpp5k-31 | Deletion | Positive? | No LoxP on target | Fail |
| Inpp5k-9 | | Negative | | Fail | Inpp5k-32 | | Negative | | Fail |
| Inpp5k-10 | | Negative | | Fail | Inpp5k-33 | Deletion | Positive | Mosiac: Possible flox | Flox |
| Inpp5k-11 | Deletion | Negative | | Fail | Inpp5k-34 | | Negative | | Fail |
| Inpp5k-12 | | Negative | | Fail | Inpp5k-35 | | Negative | | Fail |
| Inpp5k-13 | | Negative | | Fail | Inpp5k-36 | | Negative | | Fail |
| Inpp5k-14 | | Negative | | Fail | Inpp5k-37 | | Negative | | Fail |
| Inpp5k-15 | | Small band | | Deletion | Inpp5k-38 | | Negative | | Fail |
| Inpp5k-16 | | Negative | | Fail | Inpp5k-39 | | Negative | | Fail |
| Inpp5k-17 | | Negative | | Fail | Inpp5k-40 | | Negative | | Fail |
| Inpp5k-18 | | Positive | | Fail | Inpp5k-41 | | Negative | | Fail |
| Inpp5k-19 | | Negative | | Fail | Inpp5k-42 | | Negative | | Fail |
| Inpp5k-20 | Deletion | Small band | LoxP with deletion | Fail | Inpp5k-43 | | Negative | | Fail |
| Inpp5k-21 | | Negative | | Fail | Inpp5k-44 | | Negative | | Fail |
| Inpp5k-22 | | Negative | | Fail | Inpp5k-45 | Negative | Negative | ND | Fail |
| Inpp5k-23 | | Negative | | Fail | Inpp5k-46 | | Negative | | Fail |
| Inpp5k-24 | Deletion | Negative | | Deletion | Inpp5k-47 | Negative | Negative | ND | Fail |
| Inpp5k-25 | | Negative | | Fail | Inpp5k-48 | | Negative | | Fail |
| Inpp5k-26 | | Negative | | Fail | Inpp5k-49 | Negative | Negative | ND | Fail |
| Inpp5k-27 | | Negative | | Fail | Inpp5k-50 | | Negative | | Fail |

### Slide 10
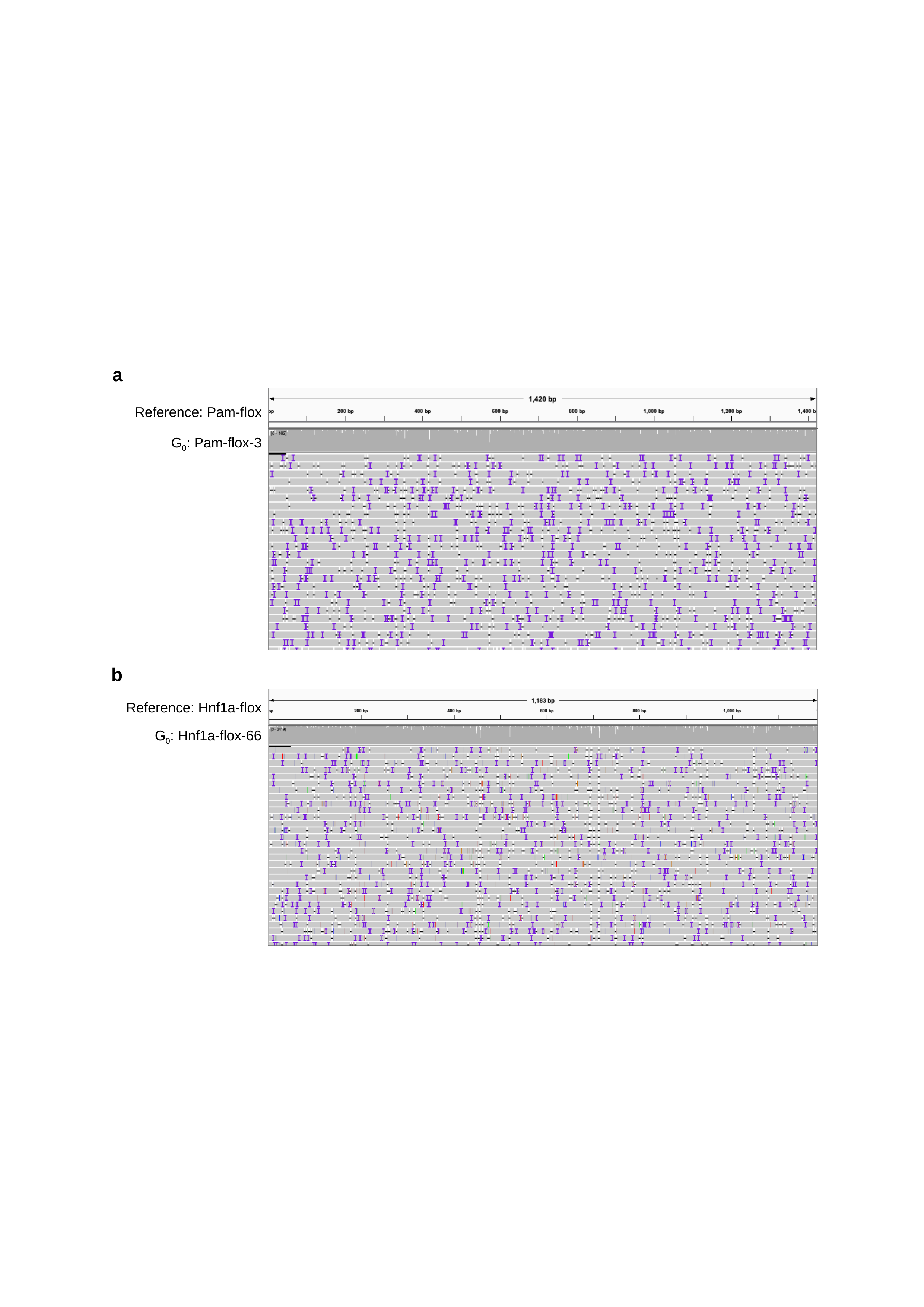

a
Reference: Pam-flox
G_0_: Pam-flox-3
b
Reference: Hnf1a-flox
G_0_: Hnf1a-flox-66

### Slide 11
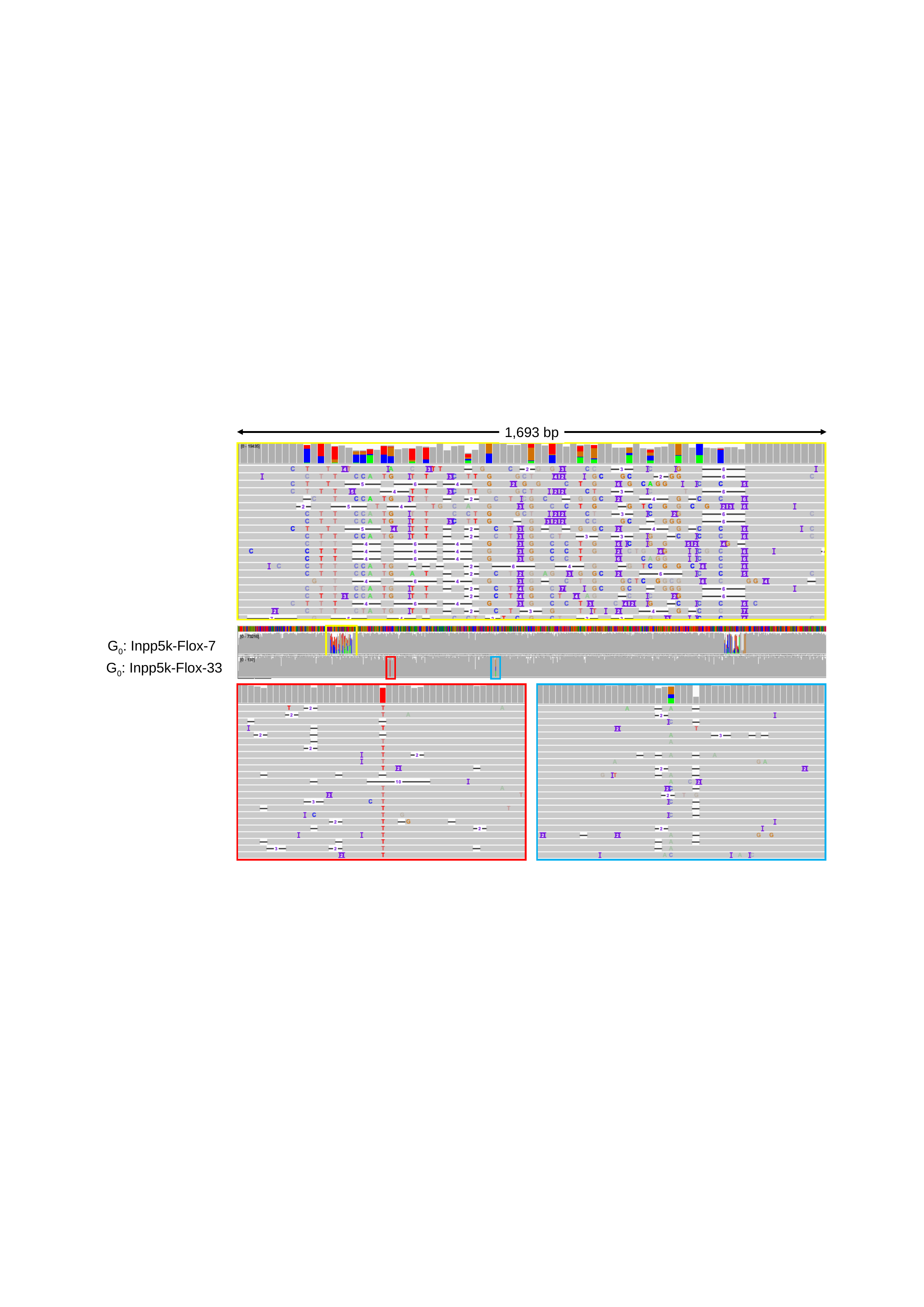

1,693 bp
G_0_: Inpp5k-Flox-7
G_0_: Inpp5k-Flox-33

### Slide 12
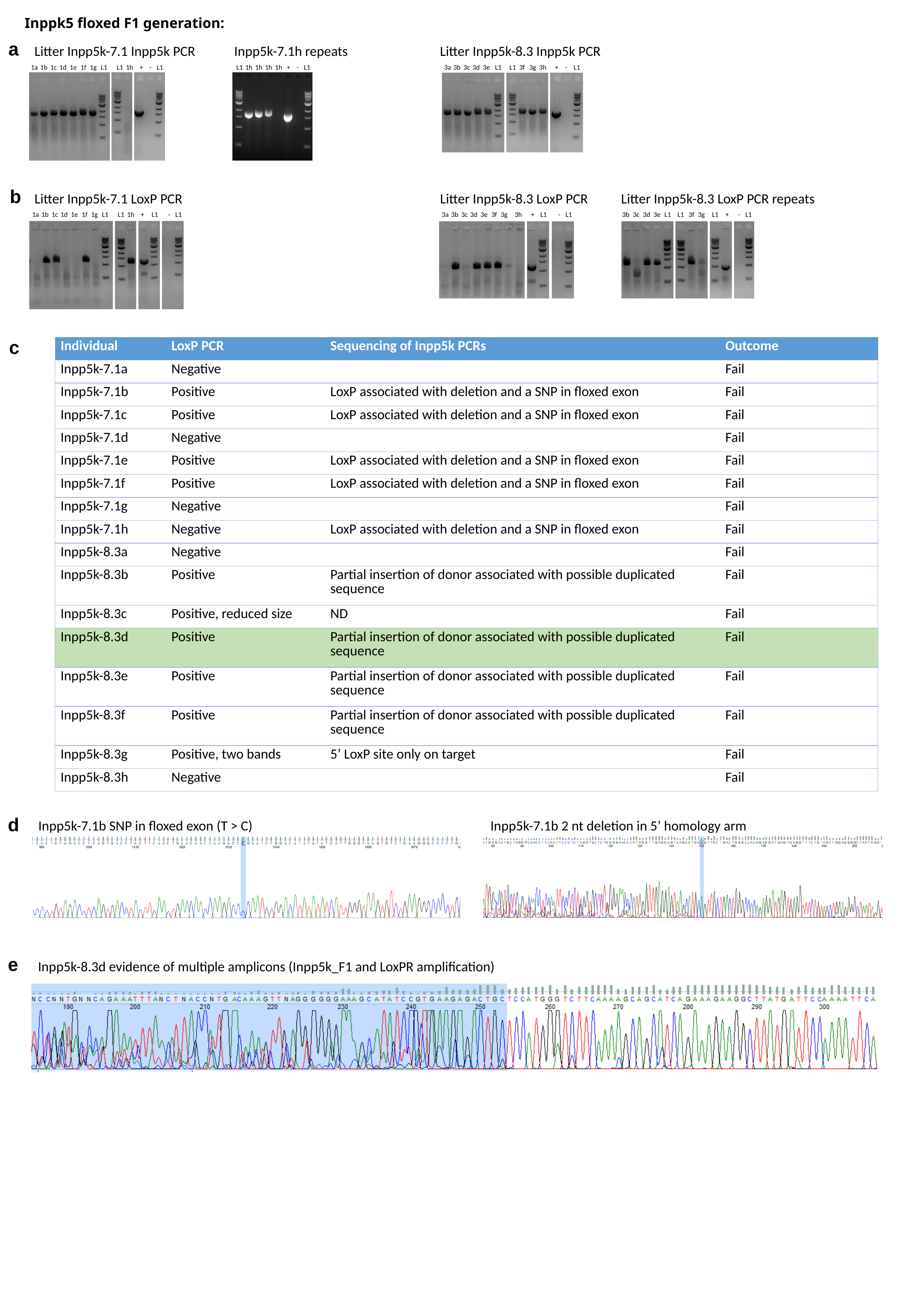

Inppk5 floxed F1 generation:
a
Inpp5k-7.1h repeats
Litter Inpp5k-8.3 Inpp5k PCR
Litter Inpp5k-7.1 Inpp5k PCR
1a
1b
1c
1d
1e
1f
1g
L1
L1
1h
+
 -
L1
L1
1h
1h
1h
1h
+
 -
L1
3a
3b
3c
3d
3e
L1
L1
3f
3g
3h
+
 -
L1
b
Litter Inpp5k-7.1 LoxP PCR
Litter Inpp5k-8.3 LoxP PCR
Litter Inpp5k-8.3 LoxP PCR repeats
1a
1b
1c
1d
1e
1f
1g
L1
L1
1h
+
L1
 -
L1
3a
3b
3c
3d
3e
3f
3g
3h
+
L1
 -
L1
3b
3c
3d
3e
L1
L1
3f
3g
L1
+
 -
L1
c
| Individual | LoxP PCR | Sequencing of Inpp5k PCRs | Outcome |
| --- | --- | --- | --- |
| Inpp5k-7.1a | Negative | | Fail |
| Inpp5k-7.1b | Positive | LoxP associated with deletion and a SNP in floxed exon | Fail |
| Inpp5k-7.1c | Positive | LoxP associated with deletion and a SNP in floxed exon | Fail |
| Inpp5k-7.1d | Negative | | Fail |
| Inpp5k-7.1e | Positive | LoxP associated with deletion and a SNP in floxed exon | Fail |
| Inpp5k-7.1f | Positive | LoxP associated with deletion and a SNP in floxed exon | Fail |
| Inpp5k-7.1g | Negative | | Fail |
| Inpp5k-7.1h | Negative | LoxP associated with deletion and a SNP in floxed exon | Fail |
| Inpp5k-8.3a | Negative | | Fail |
| Inpp5k-8.3b | Positive | Partial insertion of donor associated with possible duplicated sequence | Fail |
| Inpp5k-8.3c | Positive, reduced size | ND | Fail |
| Inpp5k-8.3d | Positive | Partial insertion of donor associated with possible duplicated sequence | Fail |
| Inpp5k-8.3e | Positive | Partial insertion of donor associated with possible duplicated sequence | Fail |
| Inpp5k-8.3f | Positive | Partial insertion of donor associated with possible duplicated sequence | Fail |
| Inpp5k-8.3g | Positive, two bands | 5’ LoxP site only on target | Fail |
| Inpp5k-8.3h | Negative | | Fail |
d
Inpp5k-7.1b SNP in floxed exon (T > C)
Inpp5k-7.1b 2 nt deletion in 5’ homology arm
e
Inpp5k-8.3d evidence of multiple amplicons (Inpp5k_F1 and LoxPR amplification)

### Slide 13
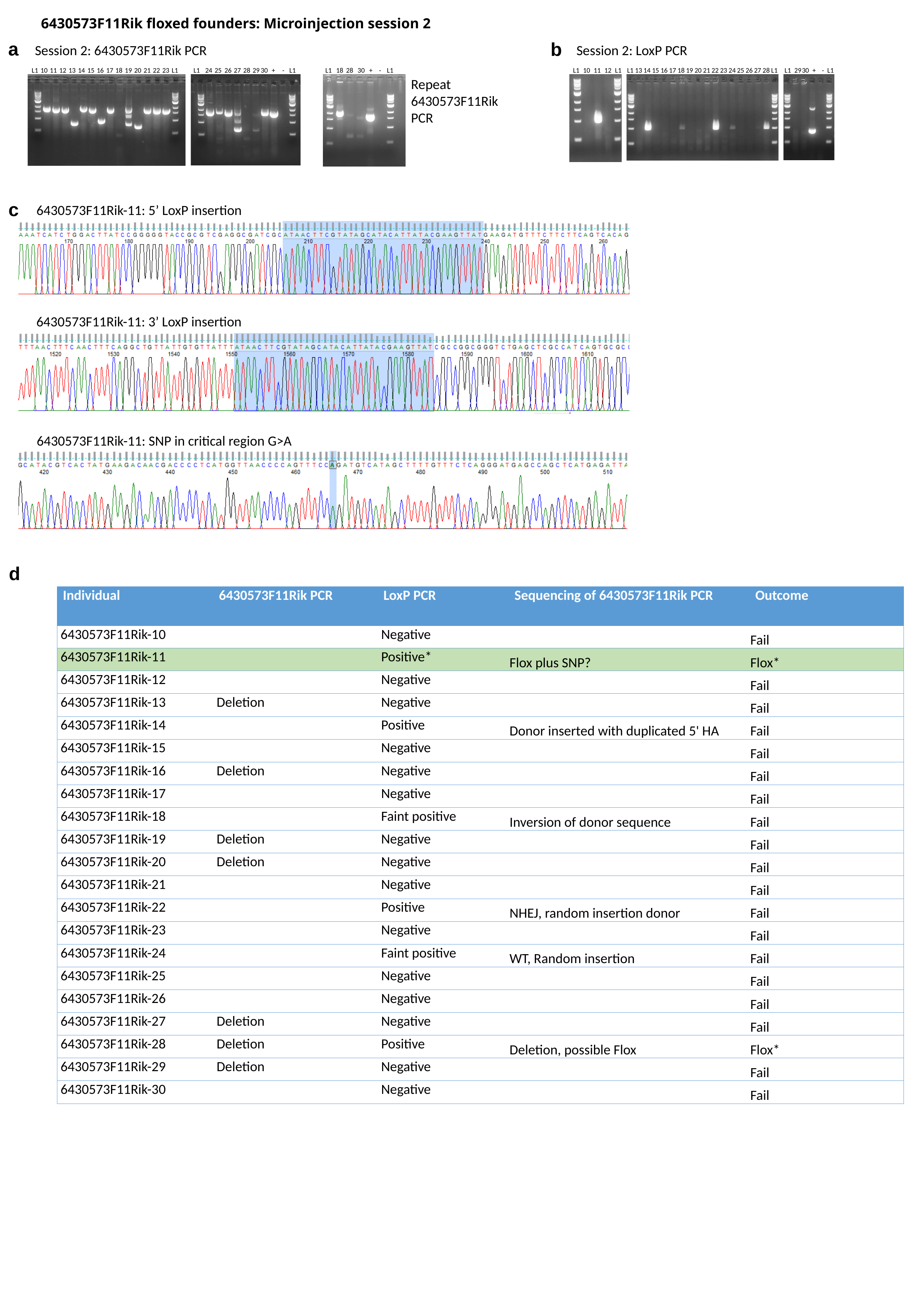

6430573F11Rik floxed founders: Microinjection session 2
a
b
Session 2: LoxP PCR
Session 2: 6430573F11Rik PCR
L1
10
11
12
13
14
15
16
17
18
19
20
21
22
23
L1
L1
24
25
26
27
28
29
30
+
-
L1
L1
18
28
30
+
-
L1
Repeat 6430573F11Rik PCR
L1
10
11
12
L1
L1
13
14
15
16
17
18
19
20
21
22
23
24
25
26
27
28
L1
L1
29
30
+
-
L1
c
6430573F11Rik-11: 5’ LoxP insertion
6430573F11Rik-11: 3’ LoxP insertion
6430573F11Rik-11: SNP in critical region G>A
d
| Individual | 6430573F11Rik PCR | LoxP PCR | Sequencing of 6430573F11Rik PCR | Outcome |
| --- | --- | --- | --- | --- |
| 6430573F11Rik-10 | | Negative | | Fail |
| 6430573F11Rik-11 | | Positive\* | Flox plus SNP? | Flox\* |
| 6430573F11Rik-12 | | Negative | | Fail |
| 6430573F11Rik-13 | Deletion | Negative | | Fail |
| 6430573F11Rik-14 | | Positive | Donor inserted with duplicated 5' HA | Fail |
| 6430573F11Rik-15 | | Negative | | Fail |
| 6430573F11Rik-16 | Deletion | Negative | | Fail |
| 6430573F11Rik-17 | | Negative | | Fail |
| 6430573F11Rik-18 | | Faint positive | Inversion of donor sequence | Fail |
| 6430573F11Rik-19 | Deletion | Negative | | Fail |
| 6430573F11Rik-20 | Deletion | Negative | | Fail |
| 6430573F11Rik-21 | | Negative | | Fail |
| 6430573F11Rik-22 | | Positive | NHEJ, random insertion donor | Fail |
| 6430573F11Rik-23 | | Negative | | Fail |
| 6430573F11Rik-24 | | Faint positive | WT, Random insertion | Fail |
| 6430573F11Rik-25 | | Negative | | Fail |
| 6430573F11Rik-26 | | Negative | | Fail |
| 6430573F11Rik-27 | Deletion | Negative | | Fail |
| 6430573F11Rik-28 | Deletion | Positive | Deletion, possible Flox | Flox\* |
| 6430573F11Rik-29 | Deletion | Negative | | Fail |
| 6430573F11Rik-30 | | Negative | | Fail |
